# Predictive Models of Genome-wide Aryl Hydrocarbon Receptor DNA Binding Reveal Tissue Specific Binding Determinants

**DOI:** 10.1101/2022.05.13.491754

**Authors:** David Filipovic, Wenjie Qi, Omar Kana, Daniel Marri, Edward L. LeCluyse, Melvin E. Andersen, Suresh Cuddapah, Sudin Bhattacharya

## Abstract

**Background:** The Aryl Hydrocarbon Receptor (AhR) is an inducible transcription factor (TF) whose ligands include the environmental contaminant 2,3,7,8-tetrachlorodibenzo-*p*-dioxin (TCDD). TCDD-mediated toxicity occurs through activation of AhR and its subsequent binding to the Dioxin Response Element (DRE), comprising the DNA motif 5’-GCGTG-3’. However, AhR binding in human tissues is highly dynamic and tissue specific. Approximately 50% of all experimentally verified AhR binding sites do not contain a DRE. Additionally, most accessible DREs are not bound by AhR. Identification of tissue specific AhR binding determinants is crucial for understanding downstream gene regulation and potential adverse outcomes of AhR activation.

**Results:** We applied XGBoost, a supervised machine learning architecture, to predict the genome wide AhR binding status of DREs in open chromatin as a function of DNA sequence flanking the DRE, chromatin accessibility, histone modifications (HM), TF binding, and proximity of the DRE to gene promoters. We trained and validated our models using 5-fold cross validation to predict the binding status of DREs in AhR-activated MCF-7 breast cancer cells, primary human hepatocytes, and lymphoblastoid GM17212 cells, as well as AhR non-activated HepG2 hepatocellular carcinoma cells. Our results demonstrate highly accurate and robust models of AhR binding; and identify patterns of transcription factor binding and histone modifications predictive of AhR binding. These patterns are consistent within tissues but highly variable across tissues, which is suggestive of tissue-specific mechanisms of AhR binding.

**Conclusions:** AhR binding is driven by a complex interplay of tissue-agnostic DNA sequence flanking its binding motif and tissue-specific local chromatin context.

## Introduction

Transcriptional regulation of gene expression is one of the main mechanisms by which cellular identity, differentiation, development and response to exogenous stimuli are coordinated (Sonawane *et al*., 2017). Gene regulation is governed in large part through direct or indirect binding of transcription factors (TFs) to DNA (Todeschini, Georges and Veitia, 2014). Changes in the activity, expression, or DNA-binding ability of even a single TF can affect the expression of hundreds to thousands of genes (Warrick *et al*., 2016; Caetano *et al*., 2018). In fact, experimental removal of even a single TF binding site, e.g., through promoter bashing or targeted mutagenesis, can result in altered gene expression (Kress, Reichert and Schwarz, 1998; Ye *et al*., 2019). In addition, TF binding is highly tissue- and cell type-specific, which makes the problem of identifying TF binding sites in a tissue-specific manner exceptionally complex (Sonawane *et al*., 2017).

The task of predicting DNA binding sites for a particular TF across the entire genome, in a specific cell type or tissue, can be likened to finding the proverbial needle in a haystack. The problem is particularly acute for TFs with short core DNA binding motifs like the aryl hydrocarbon receptor (AhR), which binds to a core 5-base pair (bp) sequence, 5’-GCGTG-3’ (Denison, Fisher and Whitlock, 1988; Swanson, Chan and Bradfield, 1995). Statistically, this motif is likely to occur in the human genome millions of times. A typical chromatin immunoprecipitation followed by sequencing (ChIP-seq) binding assay for AhR, on the other hand, yields only on the order of a few hundred to a few thousand DNA regions enriched for AhR binding, with anywhere between zero and 19 AhR core motifs appearing within each bound region (Lo and Matthews, 2012; Yang *et al*., 2018). Clearly, the occurrence of the 5-bp core motif is neither sufficient nor necessary to induce AhR binding to DNA. For one, the vast majority of putative motifs would lie in nucleosome-bound, inaccessible regions of the genome. In addition, experimental evidence suggests that specific nucleotides flanking the core motif on both 5’ and 3’ ends might form an extended active AhR binding motif (Sun *et al*., 2004). The precise identity of these flanking sequences or the mechanism by which they contribute to AhR binding is not known. Furthermore, direct, functional binding to DNA can be hard to distinguish from indirect binding through tethering with other TFs or through 3D looping of chromatin (Liang *et al*., 2014). Thus, genomic and potentially epigenomic features beyond just the DNA sequence of the core DRE motif likely play a role in determining AhR binding in vivo.

The AhR is a ligand-activated transcription factor belonging to the basic-helix-loop-helix (bHLH) PER-ARNT-SIM (PAS) family (Denison and Nagy, 2003; Abel and Haarmann-Stemmann, 2010). In addition to activation by its multiple endogenous ligands (Gutiérrez-Vázquez and Quintana, 2018; Rothhammer and Quintana, 2019), AhR mediates toxic effects of environmental contaminants such as 2,3,7,8-tetrachlorodibenzo-*ρ*-dioxin (TCDD) (Mimura and Fujii-Kuriyama, 2003; Gutiérrez-Vázquez and Quintana, 2018). Prior to ligand binding and activation, AhR is sequestered in the cytoplasm by chaperone proteins including a dimer of the 90-kDa heat shock protein (HSP90) (Denis *et al*., 1988; Perdew, 1988), the AhR-interacting protein (AIP) (Carver and Bradfield, 1997; Meyer and Perdew, 1999), and the cochaperone protein p23 (Grenert *et al*., 1997). When activated, AhR translocates to the nucleus and forms a heterodimer with the AhR Nuclear Translocator (ARNT) protein (Ikuta *et al*., 1998; Ikuta, Kobayashi and Kawajiri, 2004). The AhR-ARNT heterodimer then binds to specific DNA sequences containing the consensus 5’-GCGTG-3’ core binding motif (Durrin *et al*., 1987; Dere *et al*., 2011). This binding motif has variously been termed the Aryl Hydrocarbon Response Element (AHRE), Dioxin Response Element (DRE), or Xenobiotic Response Element (XRE) (Nebert *et al*., 2000). Here we will use the nomenclature DRE to refer to potential AhR binding sites containing the consensus core motif.

The AhR regulates a suite of target genes including Cytochromes p450 1A1 (CYP1A1), 1A2 (CYP1A2) and 1B1 (CYP1B1) by binding to DREs in the proximal promoters or, potentially, distal enhancers of these genes (Beischlag *et al*., 2008; Esser, Rannug and Stockinger, 2009; Sorg, 2014). Identification of AhR binding sites is the first step in reconstructing the AhR-mediated gene regulatory network which is crucial for understanding the role of AhR in toxicity and disease, as well as its role in physiological functions such as immune response (Esser, Rannug and Stockinger, 2009), circadian rhythm (Shimba and Watabe, 2009), cell cycle progression (Marlowe and Puga, 2005), and embryonic development (Gialitakis *et al*., 2017). High throughput molecular techniques combining the enrichment of TF-DNA complexes followed by sequencing of enriched DNA fragments, such as ChIP-seq (Barski *et al*., 2007), ChIP-exo (Rhee and Pugh, 2011) and ChIP-nexus (He, Johnston and Zeitlinger, 2015) have enabled a genome-wide view of TF binding. Over time, experiments determining the genome-wide binding profiles of hundreds of TFs in multiple cell lines, primary cells, and whole tissues have been performed and made publicly available. Likewise, AhR binding has been experimentally explored in several human cell lines and primary cells. However, the determinants of tissue-specificity of AhR binding remain poorly understood.

In recent years, multiple computational approaches for genome-wide TF binding prediction have been developed. One of the most commonly used methods relies on the position weight matrix (PWM). PWMs represent a quantitative statistical description of known and experimentally confirmed binding sites for a particular TF, effectively making up the TF’s binding motif. PWMs are derived from experimental data and can be readily obtained from databases such as TRANSFAC or JASPAR, or estimated de novo (Wasserman and Sandelin, 2004; Khan *et al*., 2018). For each potential binding site, a PWM produces a quantitative score that is a sum of individual scores of each nucleotide making up the PWM and overlapping the potential binding site. Most commonly, PWMs are used to scan the genome for TF binding sites, using a previously derived optimal threshold score as the cutoff for binding prediction (Staden, 1984; Li and Stormo, 2001). These PWMs are often derived from in vitro experiments, most commonly high throughput systematic evolution of ligands by exponential enrichment (HT-SELEX) (Ogawa and Biggin, 2012), and sometimes in vivo experiments such as ChIP-seq. However, many TFs exhibit high levels of in vivo binding to DNA sequences that do not possess the in vitro or even the in vivo derived binding motif (Karimzadeh and Hoffman, 2019).

On the other hand, TFs in eukaryotes generally do not bind DNA in isolation but rather in dense, often tissue-specific, TF clusters with binding sites of multiple TFs co-occurring in close proximity (Gotea *et al*., 2010; Yan *et al*., 2013). Consequently, PWMs of co-bound TFs could potentially be used to predict the binding of a TF of interest. However, models incorporating PWMs of co-binding TFs have shown limited utility in improving model performance (Karimzadeh and Hoffman, 2019). Nevertheless, given that TFs bind in clusters and that PWMs are not necessarily representative of actual TF binding, ChIP-seq signals of co-bound TFs, as a measure of their actual binding, are likely to provide the information that PWMs cannot. In addition, interpretable machine learning models incorporating measures of co-bound TFs could provide mechanistic insights into the determinants of tissue specific binding for a TF of interest, such as the AhR.

Over time, PWM models have been extended to include features shown to be associated with TF binding such as chromatin accessibility, histone modifications, evolutionary sequence conservation, PWMs of co-bound TFs, and gene expression (Pique-Regi *et al*., 2011; Karimzadeh and Hoffman, 2019). Similarly, a wide array of statistical modeling and machine learning approaches ranging from unsupervised Bayesian mixture models (Pique-Regi *et al*., 2011) to deep learning (Keilwagen, Posch and Grau, 2019; Quang and Xie, 2019; Srivastava and Mahony, 2020) have been applied to predict tissue-specific TF binding. However, even though some of these models achieve high cross-tissue prediction performance for some TFs, they generally lack interpretability and provide little mechanistic insight into what drives tissue or cell type specific TF binding. This problem is especially acute for TFs exhibiting highly variable binding across tissues.

In addition, most computational models of TF binding to date have been developed for and tested on constitutively active TFs. Binding of inducible TFs, such as nuclear receptors or other ligand-activated TFs like the AhR, remains largely unexplored. In this study, we applied a supervised machine learning algorithm, XGBoost (Chen and Guestrin, 2016), to develop machine learning models predicting the AhR binding status of DREs in open chromatin, i.e., DRE is bound or unbound, in four cell lines and one primary cell type: two human breast cancer cell lines (MCF-7 and T-47D) (Lo and Matthews, 2012; Yang *et al*., 2018), primary human hepatocytes (this publication), human hepatocellular carcinoma cell line (HepG2) – obtained from the ENCODE project (Dunham *et al*., 2012; Davis *et al*., 2018), and lymphoblastoid cell line (GM17212) (Neavin *et al*., 2019). The cells in these experiments were treated with either TCDD, Methylcholanthrene (3-MC; an AhR ligand) or Dimethyl sulfoxide (DMSO; vehicle control) for a duration of either 45 minutes, 1 hour or 24 hours. Using these datasets and corresponding chromatin accessibility experiments, we first identified tissue-specific AhR-bound and unbound DREs in open chromatin of each tissue. Then, we developed machine learning models that predict the binding status of DREs in open chromatin for each tissue. Our results demonstrate highly accurate and robust models of within-tissue binding. We identify several TFs as predictive of AhR binding in individual tissues, such as GATA3 in MCF-7 cells, MXI1 in HepG2 cells, and SP1 in primary human hepatocytes and GM17212 cells; as well as histone modifications (HMs) – H3K4me1 and H3K4me3 in MCF-7 cells, H3K4me3 and H3K27ac in primary hepatocytes, H3K27ac in GM17212 cells. Our tissue-specific models generalize well to the prediction of AhR binding sites without DREs, demonstrating the robustness of the models. In conclusion, we demonstrate that the patterns of TFs and HMs most predictive of AhR binding are consistent within tissues but highly variable across tissues, which is suggestive of potentially different underlying tissue-specific mechanisms of AhR binding. Additionally, we show that AhR binding is driven by a complex interplay of tissue-agnostic DNA sequence flanking the DRE and tissue-specific local chromatin context. The approach used here can be adapted to other inducible TFs, such as steroid hormone and nuclear receptors.

## RESULTS

### AhR binding profile is tissue-specific

To determine the role of the core 5’-GCGTG-3’ AhR binding motif (Dioxin Response Element – DRE) and the tissue-specific determinants of AhR binding, we compared AhR binding data across several publicly available human ChIP-seq and ChIP-chip experiments, and the primary human hepatocyte ChIP-seq experiment carried out for this study. Each experiment was performed with cells of a specific cell line (MCF-7, HepG2, GM17212, T-47D) or primary cell type (primary hepatocytes). The cells in these experiments were treated with either an AhR agonist (TCDD or 3- MC) or vehicle control – Dimethyl sulfoxide (DMSO) for a duration of 45 minutes, 1 hour or 24 hours (for the full list see **Supplementary Table 1**). In developing our machine learning models we used the data from four AhR ChIP-seq binding experiments, namely 1) MCF-7 cells treated with 10 nM TCDD for 24 hours (referred to MCF-7 or MCF-7 24h), 2) primary hepatocytes treated with 1 nM TCDD for 24 hours (referred to as primary hepatocytes), 3) HepG2 cells without AhR agonist treatment (referred to as HepG2), 4) GM17212 cells treated with 1 µM 3-MC for 24 hours (referred to as GM17212). Some additional analyses were based on the remaining three AhR ChIP-seq and ChIP-chip binding experiments – 1) MCF-7 cells treated with 10 nM TCDD for 45 minutes (referred to as MCF-7 45m), 2) T-47D cells treated with 10 nM TCDD for 1 hour (referred to as T-47D TCDD), 3) T-47D cells treated with 1 µM 3-MC for 1 hour (referred to as T-47D 3-MC). AhR binding in T-47D cells was originally investigated using the ChIP-chip experiment.

By analyzing the human reference genome - version hg19, we determined that there were close to 1.6 million DREs in the human genome. We identified the intersection of these genomic DREs with the genomic locations of AhR peaks and obtained the number of DREs under each AhR peak. We observed that AhR peaks overlap either none (0-DRE peaks), exactly one (singleton peaks), or more than one (multi-DRE peaks) DREs (**Figure 1A**). To further investigate the importance of DREs in determining AhR binding, we calculated the percentage of 0-DRE, singleton, and multi-DRE AhR peaks within each AhR binding data set listed in **Table 1** (**Figure 1B**). All data sets have a similar percentage of singleton AhR peaks ranging between 22.3% to 33%. However, the liver cells (HepG2 and primary hepatocytes) exhibited a larger proportion of multi-DRE peaks, when compared to other data sets (**Figure 1B**). Interestingly, HepG2 cells were not treated with an AhR agonist, yet the binding experiment resulted in 12,164 AhR peaks. These results suggest a level of basal induction of AhR in HepG2 and potentially other cells in cell culture conditions. Next, we investigated the distributions of singleton DRE locations relative to the mid-point of their corresponding AhR peak. If these DREs are functional in determining AhR binding, we would expect them to be enriched at one central or two centrally symmetrical points, depending on the physical position of the DNA binding domain within the AhR relative to the protein’s 3D spatial configuration. In all binding experiments, the majority of singleton DREs are situated near the mid-point of the corresponding AhR peak, i.e., they are centrally enriched (**Figure 1C, Supplementary Figure 1**). For example, in MCF-7 cells (Yang *et al*., 2018), the DREs of about 50% and 80% of singleton peaks were found within 100 and 200 base pairs up-/down-stream from the midpoint of the peak, respectively (**Figure 1C**). However, in HepG2 cells, even though singleton DREs are centrally enriched, they are markedly less centrally enriched than singleton DREs in MCF-7, primary hepatocytes and GM17212 cells (**Supplementary Figure 1**).

**Figure 1.**
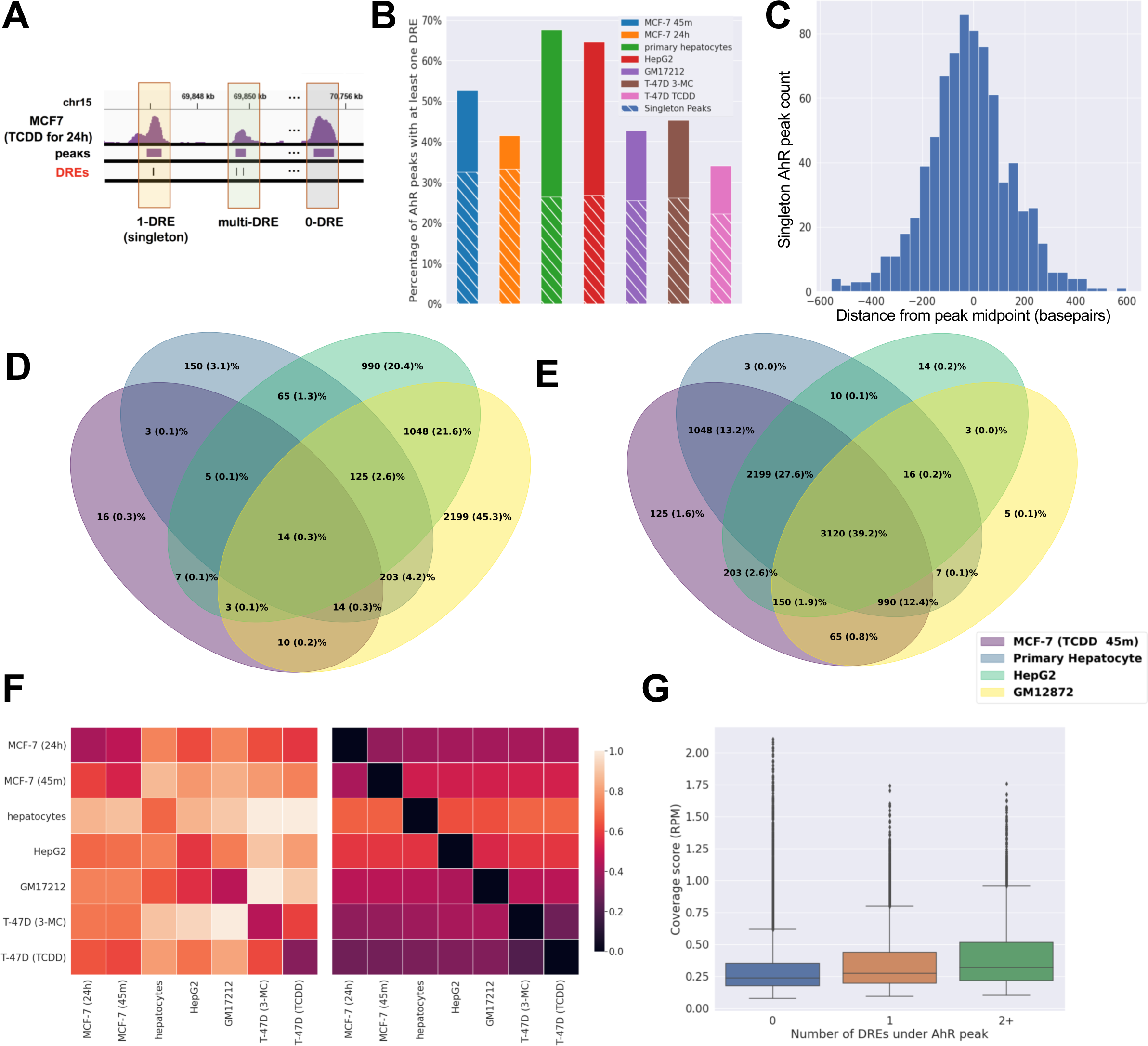
AhR binding is tissue-specific. **A** – example of AhR peaks with 1) no (0-DRE), 2) exactly one (singleton), and 3) more than 1 (multi-DRE) DREs under the peak. **B** – percentage of peaks containing at least one DRE (solid bars), and exactly one DRE (white shaded bars) per cell line with available AhR ChIP-seq data. **C** – histogram representing DRE position relative to the mid-point of singleton AhR peaks in MCF-7 cells. **D** – Venn diagram representing the overlap of bound DREs in open chromatin across four major cell lines and types. **E** – Venn diagram representing the overlap of unbound DREs in open chromatin across four major cell lines and types. **F** – heatmaps representing percentage of peaks with DREs in peaks that occur in both the column and the row cell line/type (left heatmap) and peaks that occur in the row but not in the column cell line/type (right heatmap). **G** – Boxplot of coverage scores in reads per million (RPM) for 0- DRE (blue box), singleton (orange box), multi-DRE (green box) AhR peaks in MCF-7 cells.

Cell type specificity of binding for many TFs has been shown to be influenced by chromatin accessibility – i.e., TF binding peaks most often overlap with DNase hypersensitive sites (DHS) (Dunham *et al*., 2012). To examine the contribution of chromatin accessibility in determining tissue specificity of AhR binding, we examined DNase-seq broadPeak files from the ENCODE database corresponding to the appropriate cell line or primary cell type used in the AhR binding experiment (Dunham *et al*., 2012; Davis *et al*., 2018) We identified DREs and AhR peaks appearing in open chromatin and determined that the majority of AhR peaks in all binding experiments are found in open chromatin (**Supplementary Table 2**). Due to the lack of DNase-seq or other chromatin accessibility experiments performed on cells treated with AhR agonists, all DNase-seq experiments used here represent the chromatin accessibility state prior to AhR activation (i.e., non-treated state). Further analysis revealed that the proportions and exact identities of unbound and bound DREs found in open chromatin are tissue specific. For example, out of nearly 8,000 DREs found in open chromatin across all four relevant binding experiments, about half are bound in at least one experiment, while only 14 DREs are bound in all four experiments (**Figure 1D**). On the other hand, about half of these ubiquitously accessible DREs are unbound in all four experiments (**Figure 1E**). Even when comparing two breast cancer cell lines, MCF-7 and T47-D, with similar treatment conditions, 45 minutes of 10 nM TCDD and 1 hour of 10 nM TCDD treatment, respectively, most accessible DREs are differentially bound between the two binding experiments. (**Supplementary Figure 2**). These results suggest that a DRE that lies within open chromatin of two different tissues can be bound in one and unbound in the other tissue, implying the existence of AhR binding determinants beyond the accessibility of chromatin.

On the other hand, AhR peaks that appear in common in two or more binding experiments have a higher percentage of singleton and multi-DRE peaks than AhR peaks that are unique to a single binding experiment (**Figure 1F**). Conversely, this means that AhR peaks with DREs are more likely to appear in more than one tissue than peaks with 0-DREs. Additionally, quantification of AhR ChIP-seq signal within a binding experiment revealed significant differences between average signal strength of 0-DRE, singleton, and multi-DRE peaks. The more DREs an AhR peak had, the higher the average ChIP-seq signal was under the peak. (**Figure 1G, Supplementary Figure 3**). These results jointly suggest that DREs are likely functional in determining AhR binding and that a DRE-centric approach to the investigation of tissue specificity of AhR binding could reveal important determinants and potential mechanisms driving AhR binding to DNA.

### Machine learning models accurately predict AhR binding

To better understand the tissue-specific determinants of AhR binding beyond the core DRE motif and chromatin accessibility, we developed a series of interpretable machine learning classifier models focused on the prediction of binary binding status (bound or unbound) of DREs in open chromatin. To this end we developed models with different combinations of input features for each individual binding experiment. We trained the models on DREs occurring under singleton, i.e., 1-DRE, peaks only. These DREs represent about one third of all AhR peaks, across all binding experiments. Multi-DRE peaks were not used for training the models since it is impossible to determine which specific DRE among the cluster of DREs was responsible for AhR binding. However, we used the multi-DRE peaks for model validation. We developed all our machine learning models using the gradient boosted tree algorithm of the XGBoost family of algorithms, which has been shown to handle non-linear data well (Elith, Leathwick and Hastie, 2008). These algorithms also supply metrics of feature importance, i.e., the contribution of individual input features to improving the model performance (Gregorutti, Michel and Saint-Pierre, 2017).

Our models use local chromatin features as inputs and are trained and validated to predict the binding status of singleton bound and isolated unbound DREs (see Methods), limited to DREs found in open chromatin of a particular cell line or cell type. Validation is performed using a 5-fold cross validation procedure (see Methods for details) (**Figure 2A**). The local chromatin input features that were used include

1) The DNA sequence immediately flanking the DRE. We included the flanking sequence of up to 7 nucleotides directly up- and down-stream from the DRE – previously proposed to be involved in AhR binding (Sun et al., 2004). These sequences were one-hot encoded prior to being used as model inputs.
2) Binned mean values of bigWig signals of the cell line or type most closely corresponding to the one used in the AhR binding experiment. E.g., for primary hepatocytes we used bigWig signals from experiments done in hepatocytes originated from H9, and for GM17212 we used bigWig signals from experiments done in GM12878. We used the bigWig files corresponding to i) DNase-seq (as representative of chromatin accessibility), ii) histone modification, and iii) transcription factor ChIP-seq signals from ENCODE – see methods for details (Dunham *et al*., 2012; Davis *et al*., 2018). For each bigWig signal and each DRE, we created 15 bins of width 99 base pairs. Each bin was assigned a value representing the average bigWig signal across the width of the bin. The mid-point of the central bin was positioned at the middle nucleotide of the 5-bp DRE.
3) Indicator variables of whether the DRE is found in a strict (+/- 200 bp away from a transcription start site - TSS) or loose (+/- 1500 bp away from the TSS) definition of a promoter.

**Figure 2.**
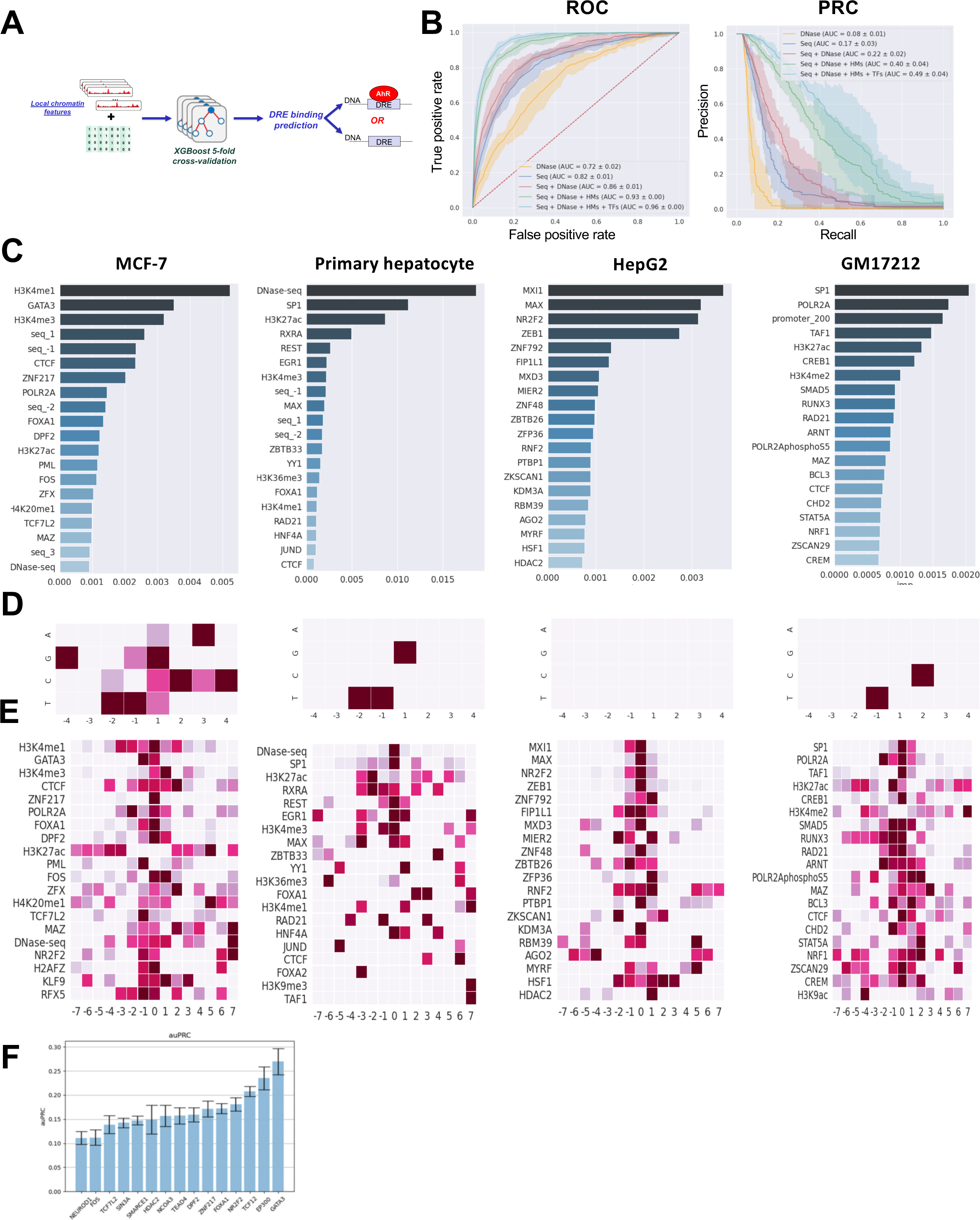
Machine learning models predicting AhR binding learn tissue specific and agnostic rules. **A** – schematic of the machine learning architecture. Local chromatin features - 1) DNA sequence flanking the DRE and 2) averaged and binned genomic signals of DNase-seq, a histone mark or a transcription factor bigWig signals are used as inputs to a XGBoost classifier which predicts the binding status of DREs in open chromatin and is trained on all accessible bound and isolated unbound DREs through 5-fold cross validation. **B** – Performance of models predicting the binding status of DREs in open chromatin of the MCF-7 cells, with five different sets of input features. Performance of each set of features is represented as a mean line with a 95% confidence interval shaded around the line resulting from 5-fold cross validation. The legend shows the list of features used, as well as area under the curve. Both receiver operating characteristic (ROC) - left panel, and precision recall (PRC) curves – right panel, are shown. **C** – Feature importance of all local chromatin context features excluding DNA sequence flanking the DRE, measured as feature importance gain in XGBoost classifier model trained on a particular cell type. Feature importance for each chromatin context feature is calculated as the average feature importance of all bins for that particular chromatin context feature. **D -** Feature importance of DNA sequence flanking the DRE, measured as feature importance gain in XGBoost classifier model trained on a particular cell line or type. Feature importance of each nucleotide type at a particular position relative to the DRE is normalized to the nucleotide type with the highest feature importance at that nucleotide position. **E** – Feature importance of all local chromatin context features excluding DNA sequence flanking the DRE in models predicting the DRE binding status of all DREs in open chromatin. Feature importance measured as feature importance gain in XGBoost classifier model trained on a particular cell line or type. Feature importance for each bin of a particular chromatin context feature is normalized to the bin with the highest feature importance for that chromatin context feature. **F –** Average area under the precision recall (PRC) curve performance across 5-fold cross validation for models predicting DRE binding status in MCF-7 cells and using only a single transcription factor as input.

To produce the best performing models, we conducted an extensive hyperparameter search for each binding experiment and input feature set (see Methods) and chose the model that produced the highest performance. In all instances, unless otherwise stated, model performance was reported as the area under the Receiver Operating Characteristic (ROC) and Precision Recall (PRC) curves, averaged over five folds using the 5-fold cross validation procedure. For class imbalanced datasets such as the ones used here, where unbound DREs far outnumber the bound (**Supplementary Table 2**), the area under the PRC curve (auPRC) is considered a more appropriate metric of model performance. Therefore, the model producing the highest auPRC was selected as the best performing model. However, the area under the ROC curve (auROC) was still a useful metric to distinguish between poorly and well performing models when comparing between binding experiments (see Methods – Performance Metrics).

To investigate model performance as a function of the input feature set, we developed and validated models with feature sets consisting of the following features - 1) DNase-seq only, 2) flanking sequence only, 3) flanking sequence and DNase-seq, 4) flanking sequence, DNase-seq and histone modifications, 5) flanking sequence, DNase-seq, histone modifications and transcription factor binding (referred to as the full model). For most cell lines and types the performance of each consecutive model improved, apart from the primary hepatocyte and HepG2 models where the performance of the sequence only model was very low overall and lower than the performance of the DNase-seq model. However, in both binding experiments the performance of the sequence and DNase-seq model is higher than the DNase-seq only model. These results suggest that even though the flanking sequence by itself provides little useful information regarding AhR binding in these binding experiments, when combined with detailed chromatin accessibility information, the flanking sequence does provide additional useful information (**Figure 2B, Supplementary Figure 4**). Additionally, most models exhibited overall high performance, except for the primary hepatocyte models (**Supplementary Figure 4**). We suspect that primary hepatocyte models did not perform as well as other models because the primary hepatocytes used in the AhR ChIP-seq experiment were obtained from a single donor specific to that experiment, while all other input features (inclusive of the list of DREs in accessible chromatin) were either obtained from a different single donor or from H9 in vitro differentiated hepatocyte-like cells (see Supplementary Materials).

To understand how specific chromatin context features contribute to the full model performance, we trained the full model on all available training data with the model hyperparameters set to values previously determined to produce the best performing model for the binding experiment of interest. Subsequently, after model training, we used the information gain metric generated by XGBoost to determine, for each binding experiment, the total feature importance of non-sequence (i.e., TF binding and epigenomic) features (**Figure 2C**), relative feature importance of sequence features per flanking sequence nucleotide position (**Figure 2D**), and relative importance of individual bins of non-sequence features (**Figure 2E**). Further examination of non-sequence feature importance scores revealed that the specific models are predominantly learning and making AhR-DRE binding predictions by relying on different features across different binding experiments (**Figure 2C**). For instance, within each binding experiment we observed three to six bigWig signals with feature importance 2-5 times higher than that of any other signal. These were: 1) in MCF-7 cells - H3K4me1, H3K4me3, GATA3, CTCF, ZNF217 and FOXA1; 2) in primary hepatocytes - DNase-seq, SP1, H3K27ac and RXRA; 3) in HepG2 cells - MXI1, MAX, NR2F2, ZEB1; and 4) in GM17212 cells - SP1, POLR2A, TAF1, H3K27ac, and CREB1. Admittedly, certain features were ranked with high importance across most binding experiments, such as the binding of CTCF, Rad21, SP1, FOXA1, MAX and MAX related factors MAZ, and MXI1; as well as H3K27ac. However, we determined that the relative level of importance of these features varied widely across different binding experiments - e.g., CTCF ranked fifth in MCF-7 cells and 20^th^ in primary hepatocytes (**Figure 2C**). Additionally, the distribution of relative importance scores across the bins were markedly different - e.g., the central bins of H3K27ac in GM17212 cells exhibited the highest importance for this feature, whereas in MCF-7 cells the central bins were not used by the model and were, consequently, not assigned an importance score (**Figure 2E**).

To investigate the contribution of DNA sequence immediately flanking the DRE, we examined the importance scores of nucleotides flanking the DRE produced by 1) flanking sequence-only models, and 2) full models (inclusive of flanking sequence, DNase-seq, histone mark and transcription factor features). We determined that the importance scores produced by the sequence only models were highly variable between binding experiments. On the other hand, the importance scores of nucleotides flanking the DRE produced by the full models revealed similar profiles of nucleotide importance across binding experiments. Mainly, the thymine residue at the flanking position directly 5’ of the DRE (labelled as the -1 position) exhibited the highest feature importance out of all nucleotides that could appear at that position in three out of four examined binding experiments (**Figure 2D**). Additionally, for two out of four binding experiments the thymine at position -2 and cytosine and guanine at position +1 also exhibited high feature importance (**Figure 2D**).

To further examine the relation of individual transcription factor binding with AhR binding within the same experiment, we developed models using only binding of a single transcription factor as input. Upon sorting these models by performance, we determined that the relative ordering of transcription factors (**Figure 2F**) was decidedly different when compared to the feature importance ordering of the full models (**Figure 2C**). Notably, for MCF-7 cells (**Figure 2F**), EP300 was the factor resulting in the second-best performing model, while in the corresponding full model, EP300 did not appear among the top 20 features with highest importance (**Figure 2C**). On the other hand, GATA3 was the most predictive factor both in the full model and individually (**Figure 2C and 2F**). These results point to a high likelihood of redundancy between binding of different TFs.

In summary, our DRE occupancy models generalized well within the binding experiment they were trained on when evaluated on a subset of bound singleton DREs and unbound isolated DREs left out from the training dataset (i.e., a single fold in a 5-fold cross validation). Additionally, these models revealed highly variable chromatin context specificity between different binding experiments, with the exception of the DNA sequence immediately flanking the DRE, where similarities between binding experiments were observed, potentially pointing to a common flanking sequence grammar.

### Singleton peak-trained models predict multi-DRE and 0-DRE AhR peak binding within the same tissue

To assess whether the high performance of our full and sequence-only models could be a result of chance due to overtraining, we performed feature selection on each of these models and compared the performance of the reduced models to the original full models. Since we use thousands of features in our full models (exact number of features depends on the cell line or cell type investigated), we created models with a subset of top N features with highest importance scores in the full model (where N = 10, 25, 50, 75, 100, 200, 300). In the MCF-7 cells we observe that the performance plateaus already at only 100 top features and that the performance of the model using only the top 50 features, although slightly lower on average, is not significantly different than the performance of the full model (**Figure 3A**). To investigate the influence of the number of flanking nucleotides included in the model we relied on a previous study that indicated, based on computational analysis of 13 experimentally verified DREs, that up to 7 up- and down-stream DRE-flanking nucleotides might play a role in determining AhR binding (Sun *et al*., 2004). However, when creating a series of models with each consecutive model including an increasing number of flanking nucleotides, we observed that the sequence-only model performance plateaus at four flanking nucleotides (**Figure 3B**). We have, therefore, used four flanking nucleotides up-/down-stream of the DRE in all our models. Additionally, since the DRE is not a palindromic binding motif and the orientation of the DRE on the genome might direct the orientation of AhR binding and its interactions with other transcription factors, we investigated whether correcting the orientation of local chromatin context features has an effect on model performance. Essentially, for DREs found on the forward strand, all features were left as-is, and for DREs found on the reverse strand, the bins of all features were flipped around the central bin (the bin containing the DRE), in order to match the DRE orientation. Our results indicate that there are no differences between our original and strand-corrected models (**Figure 3C**).

**Figure 3.**
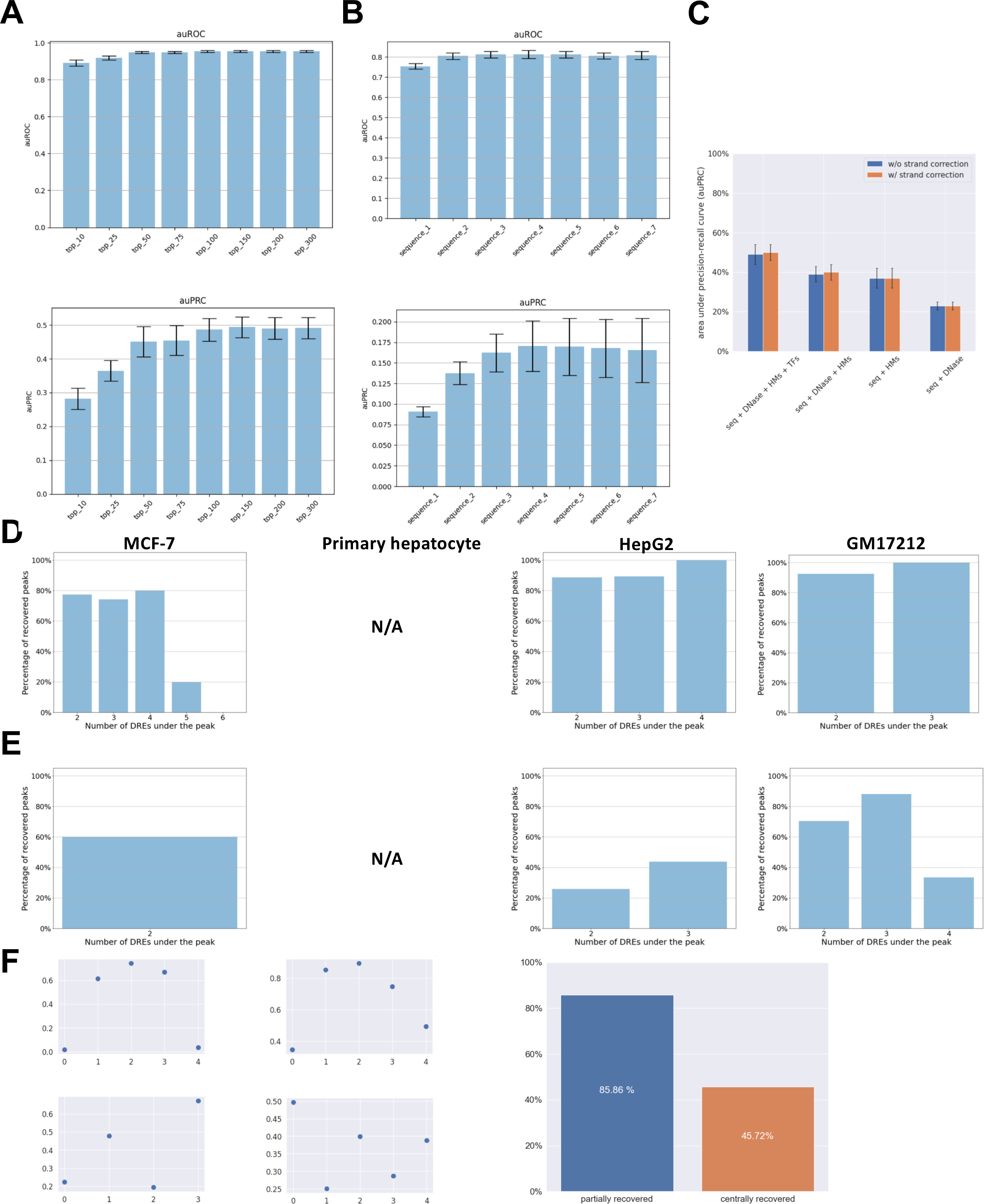
Within tissue model validation. Performance of models predicting DRE binding status in MCF-7 cells (**A, B, C and F**) measured as area under the receiver operating characteristic (top - auROC) and area under precision recall (bottom - auPRC) curve (**A and B**), **A –** containing only the top N features with highest feature performance in the full model (N=10, 25, 50, 75, 100, 200, 300) **B –** containing only the N nucleotides immediately flanking the DRE on both 5’ and 3’ sides (N=1, 2, 3, 4, 5, 6, 7). **C –** containing all features but comparing models with (blue bars) and without (orange bars) strand correction. **D –** Percentage of multi-DRE AhR peaks appearing in open chromatin, broken down by the number of DREs, that have been successfully recovered by the singleton trained model. **E** - Percentage of multi-DRE AhR peaks appearing in closed chromatin, broken down by the number of DREs, that have been successfully recovered by the singleton trained model. **F –** Examples of model output (binding probability) for four 0-DRE AhR peaks across five dummy DREs (left panel); percentage of 0-DRE AhR peaks that have been partially recovered (blue bar) and centrally recovered (orange bar)

To further examine how well the models generalize outside of their training data, we evaluated the full models for their ability to predict multi-DRE peaks. For each multi-DRE peak, we predicted the bound/unbound status of every single DRE under that peak. If at least one DRE was predicted as bound, the entire peak was considered as recovered. Recovery of multi-DRE peaks in open chromatin resulted in accuracies between 80-100% of the peaks recovered for peaks containing between two and four DREs (**Figure 3D**). Peaks containing more than four DREs were mostly not recovered by the models, suggesting a different mechanism underlying binding in areas of high DRE density, possibly through cooperative binding (Stefan and Novère, 2013). In the hg19 human reference genome approximately 1% of all DREs occur in one of these high DRE density areas, defined as 5 or more DREs within a 500-base pair region (**Supplementary Figure 5**). Similarly, multi-DRE peaks in closed chromatin are recovered at a much lower and variable rate of 25-85% (**Figure 3E**), suggesting that AhR binding in closed chromatin might be governed by a different set of rules than the binding in initially open chromatin. Contrary to the expectation that the more DREs a peak contains, the higher the likelihood of that peak being recovered by pure chance, we do not observe this trend in our results.

Similarly, we evaluated the performance of models trained on all but the DRE flanking sequence features in predicting the binding status of 0-DRE peaks. Since all non-sequence input features are calculated with the DRE genomic location as a reference and since 0-DRE peaks do not have any DREs, we simulated five “dummy DREs” for each 0-DRE AhR peak. Unlike actual DREs, these dummy DREs are not present in the genomic sequence and only represent the genomic location to be used as reference for the calculation of input features. The center of the first dummy DRE was aligned to the mid-point of the AhR peak and the other four dummy DREs were positioned at - 100, -50, +50, and +100 base pairs relative to the mid-point of the peak. We investigated up to five dummy DREs for each 0-DRE peak because the majority of DREs within singleton AhR peaks occur within –100 to +100 base pairs relative to the mid-point of the peak (as seen in **Figure 1C**). After setting up the dummy DREs we applied the same procedure as described for predicting multi-DRE peaks. Namely, a 0-DRE peak was considered partially recovered if at least one of the five dummy DREs is predicted as bound. Additionally, the peak is considered centrally recovered if the central dummy DRE is predicted as bound. In MCF-7 cells, approximately 86% of the 0-DRE peaks are partially recovered and 46% are centrally recovered (**Figure 3F**).

### Cross-tissue models provide insights into tissue-specificity of AhR binding

We developed additional models to investigate whether our full models for different binding experiments would still learn from different features even in scenarios of deliberately constrained input or output data. Specifically, we examined whether full feature models learn from different features simply because different features were available for different cell lines or cell types. To make a fair comparison between models predicting AhR binding in different cell lines or types we created models with input features limited to DNase-seq, and only those TF and HM features for which ChIP-seq experiments were available in all cell lines or types. Our results still indicate a highly variable set of the most important features determining AhR binding within different cell lines or cell types (**Figure 4A**). To further investigate whether there are enhancer-specific binding rules and whether these rules might be similar between cell lines or cell types, we created full models predicting the occupancy of singleton bound and isolated unbound DREs found only in enhancers. Once again, our results show highly variable sets of the most important features for each cell line/type. However, we observed an increase in feature importance for the EP300 transcriptional coactivator for all cell lines/types, (**Supplementary Figure 6 and Figure 2C**).

**Figure 4.**
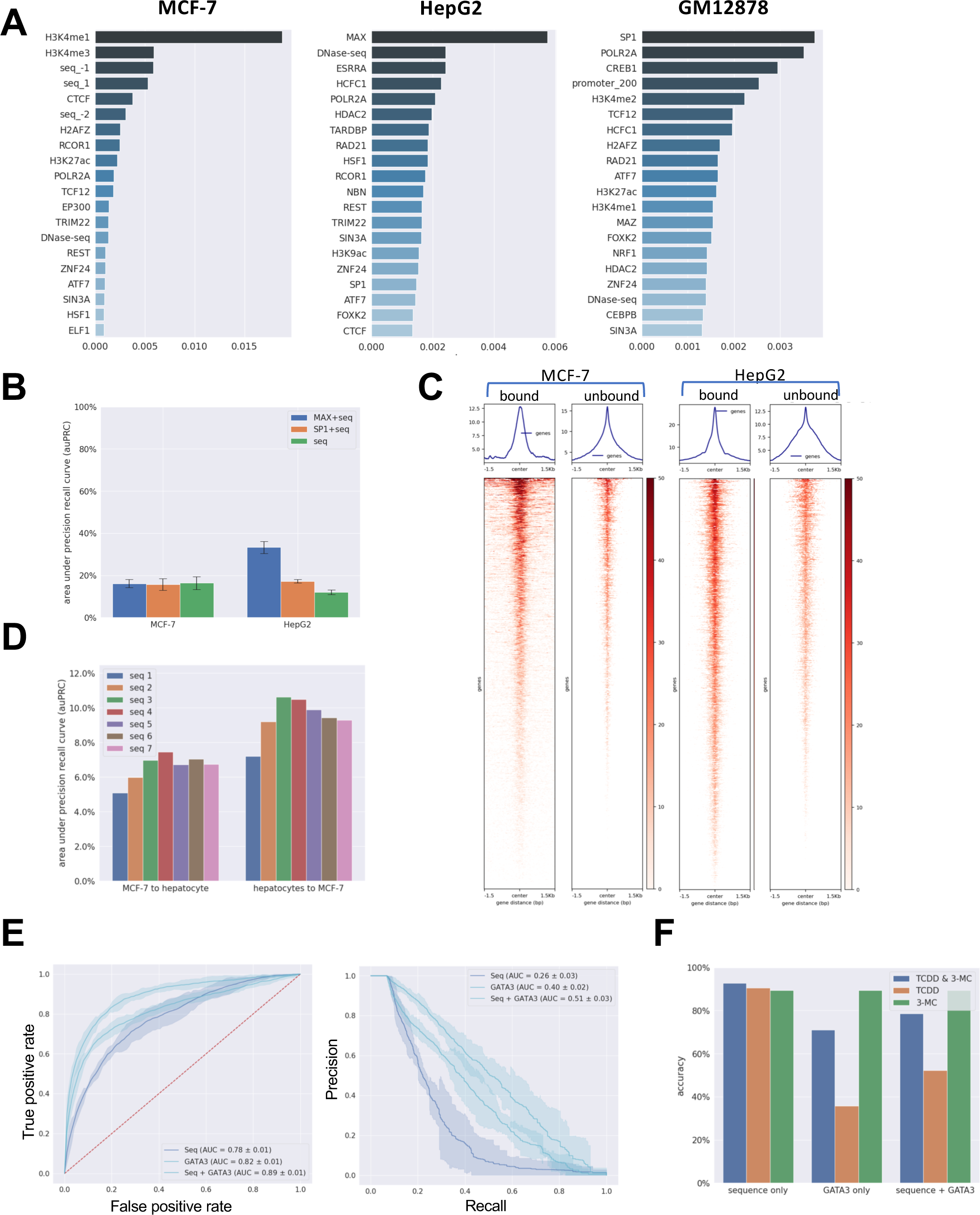
Cross tissue model validation. **A –** Feature importance of local chromatin context features excluding DNA sequence flanking the DRE, measured as feature importance gain in XGBoost classifier model trained on a particular cell line or type. Feature importance for each chromatin context feature is calculated as the average feature importance of all bins for that particular chromatin context feature. Only chromatin context features that have been ChIPed in all three cell lines were used in model training. **B –** Performance of 1) SP1 and flanking sequence model, 2) MAX and sequence only model; compared to the performance of sequence only model trained in the MCF-7 cells treated with TCDD for 24 hours. **C –** Cross tissue performance of sequence only models trained on MCF-7 cells and primary hepatocytes and evaluated on primary hepatocytes and MCF-7 tcells, respectively. **D.** Performance of models predicting the binding status of DREs in open chromatin of the MCF-7 cells, with three different sets of input features (flanking sequence only, GATA3 only, flanking sequence and GATA3). Performance of each set of features is represented as a mean line with a 95% confidence interval shaded around the line resulting from 5-fold cross validation. The legend shows the list of features used, as well as area under the curve. Both receiver operating characteristic (ROC) - left panel, and precision recall (PRC) curves – right panel, are shown. **E –** Accuracy of DRE binding prediction of the models trained in MCF-7 45m cells and evaluated in T-47D TCDD and 3-MC cells. Models are trained with 1) flanking sequence only features (left), GATA3 only (middle), flanking sequence and GATA3 (right) and evaluated on bound DREs appearing in TCDD treated cells only (orange bars), 3-MC treated cells only (green bars), both TCDD and 3-MC treated cells (blue bars).

SP1 and MAX transcription factor binding have the highest feature importance in the full feature models for GM17212 and HepG2 cells, respectively. However, in MCF-7 cells, neither of these factors appeared within the top 20 factors with highest feature importance (**Figure 2C**). To evaluate whether the feature importance of these factors within the MCF-7 cells was low due to potential redundancy with other factors, we developed MCF-7 specific models containing 1) SP1 and flanking sequence features; and 2) MAX and flanking sequence features. The average auPRC performance of these models across 5-folds was not significantly better than the performance of the corresponding sequence-only model (**Figure 4B**). These results suggest that these two factors do not play a significant role in AhR binding within the MCF-7 cell line, while they are key players in AhR binding in GM17212 and HepG2 cells. Similarly, GATA3 is the single factor most predictive of AhR binding in MCF-7 cells; however, GATA3 is either very lowly or not expressed at all in other cell types, making it hard to investigate whether the AhR binding dependence on GATA3 in MCF-7 cells is specific to that cell line. To further understand why including certain transcription factor information as model inputs results in high performing models in one tissue but not in another, we focused on the case of the transcription factor MAX. As shown in **Figure 4B**, MAX is mostly ignored by the MCF-7 model, while it is one of the most important model inputs in the HepG2 model. When comparing the binding profiles of MAX centered on bound and unbound DREs, we observed that the difference in MAX signal between bound and unbound DREs is less pronounced in MCF-7 cells than it is in HepG2 cells, which could explain the increased utility of MAX in the HepG2 model. **(Figure 4C).**

To investigate whether models specific to one cell line/type can predict AhR binding in others, we evaluated the cross-cell performance of a few models. The sequence-only model trained on MCF-7 cells and evaluated on primary hepatocytes did not perform any better than random (**Figure 4D**). Even though the cells in these two binding experiments were both treated with TCDD for 24 hours, the concentration of TCDD was different: 10 nM and 1 nM, respectively. Interestingly, the performance of the sequence-only model trained on primary hepatocytes was lower when evaluated within primary hepatocytes than when evaluated cross-tissue in MCF-7 cells (**Figure 4D**). These results indicate that cells with a relatively smaller number of bound and unbound DREs in open chromatin (such as MCF-7) might result in somewhat overfitted sequence models, especially when a large number of flanking nucleotides is used in the model.

On the other hand, 1) sequence-only, 2) GATA3-only, and 3) sequence and GATA3 models trained on the MCF-7 cells exhibit high performance within tissue, with model performance increasing with each successive model (**Figure 4E**). However, when we evaluated these models on all bound DREs within T-47D cells treated with either 3-MC or TCDD for one hour (after excluding any DREs that were bound in both MCF-7 and T-47D cells, to avoid using training data in the testing phase), it was the sequence-only model that produced the highest accuracy of 90.67%. Performance of the sequence-only model trained on MCF-7 cells evaluated separately in T-47D cells for three groups of AhR peaks appearing in 1) TCDD treatment only, 2) 3-MC treatment only, 3) and both TCDD and 3-MC treatment, yielded accuracies of 90.48%, 89.47%, and 92.86%, respectively, suggesting that the sequence-only model was unable to distinguish between AhR binding resulting from activation by different AhR ligands (**Figure 4F**). However, this result also suggests that AhR peaks in T-47D cells common to both ligands were slightly more predictable by the MCF-7 sequence-only model. These results jointly suggest that cross tissue model predictions might be more accurate for binding experiments with similar cell lines/types (like MCF-7 and T-47D, both of which are breast carcinoma cell lines) and treatments, as opposed to binding experiments with dissimilar cell types (like MCF-7 and primary hepatocytes) but similar treatments. On the other hand, the robustness of the primary hepatocyte sequence-only cross-tissue results suggest the existence of a DRE flanking nucleotide grammar that is preserved across different cell lines/types and treatments.

### AhR binding models reveal positive and negative regulators of AhR binding

To examine the decisions the AhR binding models make when predicting the binding status of each DRE, we used ELI5 (Korobov and Lopuhin, 2020), an algorithm that summarizes the decision-making process underlying individual model predictions. For each prediction, ELI5 assigns a numerical weight to each feature the model used to make that prediction. These weights represent a measure of how much a particular feature contributed to the final prediction across all decision trees used by the XGBoost model. The higher the assigned weight the more the feature contributed to the model making the exact prediction. An example of the top ten positively and negatively weighted features used in a single DRE binding status prediction in the HepG2 model is shown in **Figure 5A**. In this example, the high average bigWig signal value within bin 0 of MAX binding is assigned the highest weight and contributes most to the model predicting the corresponding DRE as bound.

**Figure 5.**
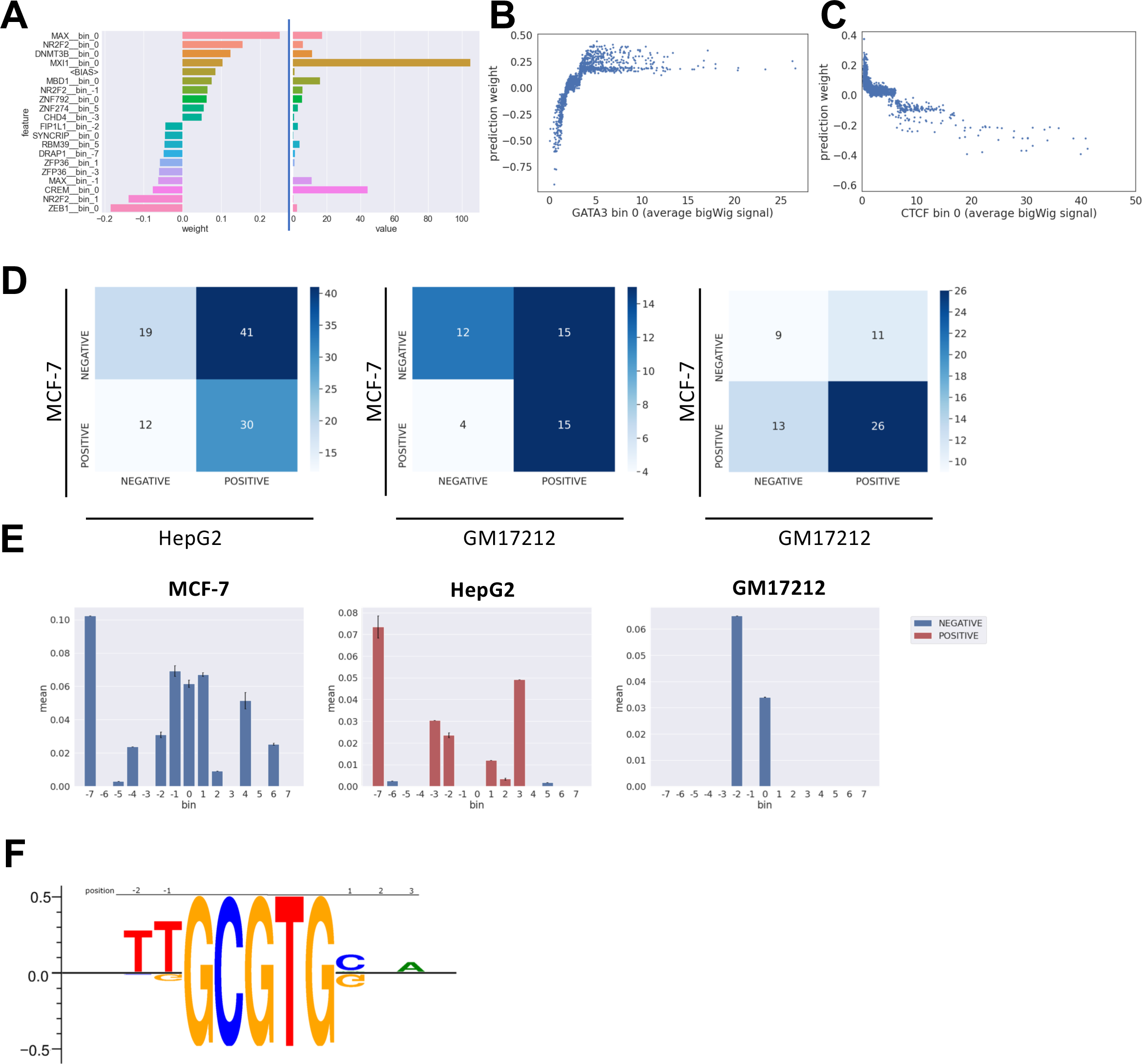
Positive and negative regulators of AhR binding. **A –** feature importance, i.e., weights (left) and feature values, i.e., average bigWig signal within the feature bin (right) of the top 10 most and least important features in the prediction of the binding status of a single bound DRE in HepG2 cells. **B –** example of bin 0 of GATA3 as a positive regulator in MCF-7 cells. The weights increase with increasing GATA3 signal. **C –** example of bin 0 of CTCF as a negative regulator in MCF-7 cells. The weights decrease with increasing CTCF signal. **D –** confusion matrices providing an overview of the number of positive and negative regulators for each pair of cell lines. **E –** average feature importance of each bin of CTCF in three investigated cell lines. Bins without a bar represent features that were not used by the model. Bars colored blue or red represent positive or negative regulators. **F –** sequence logo for binding of AhR in MCF-7 cells derived from feature importance of each nucleotide type at each flanking position, relative to the DRE. Nucleotides with positive importance contribute to the model predicting DREs as bound and nucleotides with negative importance contribute to the model predicting DREs as unbound.

First, we analyzed the weights assigned to features that are the individual bins of bigWig signals originating from DNase-seq, histone modification or transcription factor ChIP-seq experiments. When examining the weights assigned to a single feature, for example bin 0 of GATA3 binding, across all correctly classified bound DREs (true positives – TPs), we observed model features whose weights increase with increasing feature values - positive regulators (**Figure 5B**), as well as features whose weights decrease with increasing feature values - negative regulators (**Figure 5C**). To figure out whether the direction of regulation (positive or negative) of individual features is preserved between models, we investigated features in common to each combination of two models (excluding primary hepatocytes). In total we found 102, 46, and 59 features used in common by 1) MCF-7 and HepG2, 2) MCF-7 and GM17212, and 3) HepG2 and GM17212 models, respectively. For each model combination approximately 41-62% of features are positive regulators in one model and negative in another **(Figure 5D**). Additionally, out of 15 features used by all three models, only three features have the same direction of regulation in all three models – bin 1 of MAZ, bin 1 of MAX, bin 3 of H3K27ac. All three of these features are positive regulators in all three models (results not shown). Even for a single transcription factor, such as CTCF in HepG2 cells, some bins can be positive regulators, while others can be negative regulators or not be used by the model at all **(Figure 5E**). These results suggest that even though tissue-specific models primarily learn from entirely different features, a small subset of those features shows similar patterns across tissues.

Our models also provide weights for the DRE-flanking DNA sequence features. These features are binary and indicate whether a certain nucleotide is found at a certain position relative to the DRE, e.g., whether the nucleotide at position –1 is a thymine or not. These flanking sequence features can also be seen as positive or negative regulators, as the model produces either positive or negative weights when a nucleotide is present. The classical representation of sequence features across many binding sites is the sequence logo (Schneider and Stephens, 1990). These motifs indicate how informative the presence of a certain nucleotide at a particular position is in determining whether the sequence represents a putative binding site. The logo generated by our models is a combination of two motifs – one that describes bound DREs, and another that describes the unbound DREs (the upper and lower motifs in **Figure 5F,** respectively). Additionally, when comparing the direction of regulation for DNA flanking sequence features between MCF-7 and primary hepatocytes, all three nucleotide features (position -2 is thymine, -1 is thymine and 1 is guanine) that are used by both models have the same direction of regulation (results not shown).

## DISCUSSION

Even though binding of transcription factors (TFs) to DNA has been extensively studied, and many predictive models have been developed for TF binding prediction (Karimzadeh and Hoffman, 2019; Keilwagen, Posch and Grau, 2019; Quang and Xie, 2019), the determinants and mechanisms underlying tissue-specific TF binding are not well understood. In addition, unlike many constitutively active TFs, tissue-specific binding of AhR, a ligand-inducible TF, cannot be fully determined through chromatin accessibility, the extended binding motif of AhR, the motifs of other co-bound TFs, or any combination of these features.

Traditionally, ligand-activated AhR has been assumed to bind to the 5’-GCGTG-3’ DNA sequence, known as the dioxin response element (DRE), However, in vivo experimental studies have shown that DREs are not necessary for AhR binding (Lo and Matthews, 2012; Yang *et al*., 2018). The fact that AhR binds to many sequences without DREs, brings into question whether AhR-bound sequences containing DREs represent direct binding of AhR to those DREs. Our results show that 1) the number of DREs under AhR peaks relative to peak widths is higher than expected by chance, especially for peaks with multiple DREs (Figure 1B). Secondly, DREs occurring under singleton AhR peaks are centrally enriched relative to the mid-point of their respective peaks (Figure 1C), and AhR peaks with a higher number of DREs under the peak have a statistically higher average bigWig signal than AhR peaks with a lower number of DREs (Figure 1G). These results suggest that DREs, if not necessary, are functional in determining the strength of AhR binding. Interestingly, our singleton DRE-centered models of AhR binding were able to predict the binding of 0-DRE AhR peaks with high accuracy.

One possible explanation of these results is that there is a high level of indirect binding of AhR within singleton and 0-DRE AhR peaks, possibly due to tethering to other TFs that are directly bound to DNA (Lonard and O’Malley, 2006). If the number of indirectly bound DREs used in the model training phase was high enough to capture sufficient information about tethering complexes or chromatin context surrounding such complexes, then the model would be learning how to identify indirect AhR binding. The model could, therefore, predict the binding of 0-DRE peaks, which are most likely indirectly bound, with high accuracy. This explanation would also account for why singleton AhR models perform poorly in predicting the binding of AhR peaks with five or more DREs (Figure 3D), since these peaks are more likely to be bound by AhR directly and possibly also cooperatively.

Another likely explanation is that some of the bound AhR might be part of chromatin loops, with AhR being directly bound to one or more DREs, in at least one loop anchor, and the imprint of AhR binding appearing in other anchors due to their proximity. Our results show that EP300 is one of the factors most predictive of AhR binding in enhancers **(Supplementary Figure 6).** EP300 has been implicated as a participant in the formation of chromatin loops, specifically as a marker of pre-established enhancer anchors that appear in enhancer-promoter loops formed after glucocorticoid receptor (GR) activation (McDowell *et al*., 2018). Additionally, EP300, H3K4me1, and H3K27ac have been shown to jointly be markers of active enhancers (Creyghton *et al*., 2010). In this scenario, the directly bound AhR molecule could leave an impression of a 0-DRE AhR peak in another anchor of the same loop, due to their physical proximity (Liang *et al*., 2014). Any TF or chromatin complexes established in the formation of these loops would then be associated with both singleton and 0-DRE peaks. In this case our models would be learning how to identify direct AhR binding.

Most likely, AhR peaks are a result of a balance of both direct and indirect binding and in cases where this balance is heavily tipped towards indirect binding, machine learning models based on bound and unbound DREs might not be the best approach to study the binding of AhR. This could be the case for the HepG2 binding experiment where singleton DREs are not as centrally enriched as they are in other binding experiments. Unfortunately, there is currently no way of differentiating between direct and indirect binding using only ChIP-seq signals.

Machine learning models predicting transcription factor binding often investigate the possibility and performance of cross-tissue predictions, but seldom look into the extent of tissue specificity or why these models fail to perform as well as they do within-tissue. We found that AhR binding is highly tissue-specific, with only five AhR peaks appearing in common across five examined human cell lines and types, out of a total of about 30,000 distinct AhR peaks. Our results demonstrated that binding of AhR is partially determined by 1) a common cross-tissue flanking-sequence syntax (Figures 2D and 5G), and 2) tissue-specific chromatin context syntax (Figures 2C, 4A, 4B). Interestingly, the chromatin context determinants differ vastly between tissues (Figure 2C, 2E and 5D), with one or two distinct TFs dominating as the most important contributors to model performance in each model. We propose that these factors either 1) play a major role in determining cell identity, or 2) are a common AhR co-factor in that specific tissue. As an example, GATA3 is mutated in MCF-7 cells, but not in T-47D cells. The MCF-7 mutation of GATA3 is heterozygous, and results in a copy of the GATA3 protein that is more stable and resistant to turnover, and binds to DNA more strongly than its wildtype counterpart (Adomas *et al*., 2014; Takaku *et al*., 2020). We propose that it is precisely this increase in binding activity of GATA3 in MCF-7 cells that makes GATA3 the most predictive factor of AhR binding in those cells. In addition, GATA3 possesses pioneering factor activity and was also shown to be the most commonly overlapping factor for binding of ERα, another inducible TF, in MCF-7 cells (Jiang *et al*., 2019). On the other hand, AhR peaks in T-47D cells are still correlated with GATA3 binding but to a lesser extent. Surprisingly, in MCF-7 cells it was TCDD-activated AhR that was highly GATA3-dependent, while in T-47D it was 3-MC activated AhR. These results indicate that wild type GATA3 might still play a role in determining AhR binding; However, this role is not as pronounced, and is manifested in a ligand dependent manner. Since AhR binding in T-47D cells was investigated with the ChIP-chip assay, these experiments produced only a limited number of AhR enriched regions, mostly in gene promoters. Therefore, it remains unclear whether 3-MC promotes GATA3-dependent AhR binding in T-47D cells genome-wide, or only in a subset of AhR-enriched regions. Additionally, synergistic effects of GATA3 and AhR binding on the expression of GPR15 have been reported in human CD4+ T cells (Swaminathan *et al*., 2021), suggesting that AhR-GATA3 interactions might not be confined to breast cancer cells.

In contrast to pioneering TFs like GATA3, certain TFs known to be functionally associated with AhR have not been ranked highly by our models. One such TF is ARNT, which is the prototypical dimerization partner of AhR. Cobinding with ARNT is thought to account for the majority, if not all, of TCDD-induced AhR bound sites. However, our HepG2 and GM17212 models rank ARNT features very lowly (Figure 2C). Admittedly, for the GM17212 model, the ARNT binding experiment comes from a highly similar, but different tissue compared to the AhR experiment (GM12878). Both are lymphoblastoid cell lines but have been generated from different donors. Additionally, the ARNT binding experiment has no AhR ligand treatment, while the AhR binding experiment is based on treatment with 3-MC. All these factors might contribute to the observed discrepancy between AhR and ARNT binding signals, which are generally expected to be highly similar to each other. On the other hand, HepG2 cells were not treated with an AhR ligand. However, a photoproduct of tryptophan which is present in cell culture media has been shown to be an AhR agonist and could explain the basal AhR activity in HepG2 cells (Öberg *et al*., 2005). In this case the ARNT experiment would have received the same treatment as the AhR experiment. However, ARNT features were not used by the HepG2 model at all. We suspect that the tryptophan derivative-induced activation of AhR is similar to endogenous activation of AhR and does not result in high levels of AhR-ARNT dimerization. Both AhR and ARNT have biological activities, distinct from and likely equally or more important than those associated with the gene expression regulated by the AHR-ARNT heterodimer responding to exogenous ligands. Some of these activities are likely mediated through AhR di- and multi-merization with its other known partners, such as KLF6, RELA or RB1. For instance, AhR forms a multimeric complex, including retinoblastoma (RB1), CDK4 and E2F1, leading to RB1 hyperphosphorylation, facilitating cell cycle progression (Barhoover *et al*., 2010). These dimers or multimers seem to bind to DNA motifs distinct from DREs (Wilson, Joshi and Elferink, 2013). Surprisingly, even though KLF6 and RELA features were used in the HepG2 model, they did not rank highly in feature importance. However, this result might be due to the fact that we had used a DRE-centered approach and AhR-KLF6 dimers do not appear to directly bind DREs, making our selection of unbound DREs in open chromatin of HepG2 cells non-informative for the machine learning models.

It has been suggested that the DNA sequence immediately flanking the DRE forms an extended AhR binding motif and thus influences its binding (Sun *et al*., 2004). Even though our models do suggest the existence of a common flanking sequence grammar across tissues, flanking sequence features have higher feature weights in MCF-7 cells than in any other tissue. They are also used more often in individual DRE binding status predictions in MCF-7 cells than any other tissue. Therefore, it appears that certain AhR binding sites rely more on the DRE flanking sequence than others. Additionally, the number of these flanking sequence-dependent sites varies across tissues.

In summary, we have developed highly accurate and robust predictive models of within-tissue AhR binding for several distinct cell lines and one primary cell type. Our models dissected the tissue specificity of AhR binding and showed that tissue-specific AhR binding is driven by a complex interplay of tissue-agnostic DNA sequence immediately flanking the DRE, and a highly tissue-specific local chromatin context.

## Methods

### Reference genome

Unless otherwise specified the reference genome used for sequence alignment was the human genome assembly version hg19. We opted for hg19 due to availability of data on ChIP-seq and DNase-seq data repositories such as Geo Datasets (Barrett *et al*., 2013), ChIP-Atlas (Oki *et al*., 2018) and ENCODE (Dunham *et al*., 2012; Davis *et al*., 2018). Likewise, most other transcription factor binding prediction tools available at this time were trained on hg19.

### Cell culture and treatment

Primary human cryopreserved hepatocytes were obtained from Triangle Research Labs (Research Triangle Park, NC). Cells were plated on Rat Collagen I-coated plates and Geltrex was added for a final concentration of 0.25 mg/ml approximately 6 h after plating. Cells were maintained at 37 °C in a humidified 5% CO2 atmosphere in William E medium supplemented with cell maintenance cocktail B (Invitrogen). TCDD was obtained from Dow Chemicals. The cells were treated with a final concentration of 1 nM TCDD in 0.1% DMSO. and DMSO (0.1%) was used as vehicle control.

### Chromatin immunoprecipitation

Human hepatocytes incubated with 1 nM TCDD or DMSO were fixed with 1% formaldehyde at room temperature for 15 min and quenched with 0.125 M glycine and shipped to Active Motif (Carlsbad, CA, U.S.A) for ChIP-seq analysis. Chromatin was isolated by adding lysis buffer, followed by disruption with a Dounce homogenizer. Lysates were sonicated using a Misonix Sonicator 3000 equipped with a microtip in order to shear the DNA to an average length of 300– 500 bp. Lysates were cleared by centrifugation and stored at −80 °C.

Genomic DNA was prepared by treating aliquots of chromatin with RNase, proteinase K and heat for de-crosslinking, followed by phenol/chloroform extraction and ethanol precipitation. Purified DNA was quantified on a NanoDrop spectrophotometer. Extrapolation to the original chromatin volume allowed quantification of the total chromatin yield.

For each ChIP reaction, 30 µg of chromatin was precleared with protein A agarose beads (Invitrogen). ChIP reactions were set up using precleared chromatin and a rabbit polyclonal antibody against AhR (ENZO Lifesciences, BML-SA210, Lot# 01031447) and incubated overnight at 4 °C. Protein A agarose beads were added and incubation at 4 °C was continued for another 3 h. Immune complexes were washed two times each with a series of buffers consisting of the deoxycholate sonication buffer, high salt buffer, LiCl buffer, and TE buffer. Immune complexes were eluted from the beads with SDS buffer and subjected to RNase treatment and proteinase K treatment. Crosslinks were reversed by incubation overnight at 65 °C, and ChIP DNA was purified by phenol-chloroform extraction and ethanol precipitation.

### ChIP sequencing

ChIP and Input DNAs were prepared for amplification by converting overhangs into phosphorylated blunt ends and adding an adenine to the 3′-ends. Illumina genomic adapters were ligated and the sample was size-fractionated (200–250 bp) on a 2% agarose gel. After a final PCR amplification step (18 cycles), the resulting DNA libraries were quantified and sequenced on HiSeq 2000. Sequences (50 nucleotide reads, single end) were aligned to the human genome (hg19) using the BWA algorithm (Li and Durbin, 2009). Aligns were extended in silico at their 3′- ends to a length of 150 bp, which is the average genomic fragment length in the size-selected library and assigned to 32-nt bins along the genome. Peak locations were determined using the MACS algorithm (v1.4.2) with a cutoff of p = 10^−7^ (Zhang *et al*., 2008). Signal maps and peak locations were used as input data to Active Motif’s proprietary analysis program, which creates excel tables containing detailed information on sample comparison, peak metrics, peak locations and gene annotations.

### Visualization of ChIP-Seq Signal

Bigwig files were used as inputs to deepTools version 3.5.1 (Ramírez *et al*., 2014) for visualization. DeepTools heatmap function was used to create visualizations of ChIP-Seq signal fold enrichment within a −1.5 to +1.5 kb region around the bound and unbound dioxin response elements (DREs), as well as to generate average profiles for ChIP-Seq enrichment in the same region.

### DREs in open chromatin

We obtained DNase-seq experiments for all relevant cell lines (MCF-7, T-47D, primary hepatocytes, HepG2, GM12878) from ENCODE https://encodeproject.org/. We downloaded the broadPeak DNase-seq files for the hg19 genome assembly, and if there were multiple replicates, found the intersection of all replicates. Any DRE found under the peaks of DNase-seq intersection was considered to be in the open chromatin of the corresponding cell line and was used in the determination of bound and unbound DREs for the purposes of model training. DREs occurring in blacklisted regions were ignored in downstream analyses.

### AhR-bound and unbound DREs

Firstly, we assembled a list of all DREs in the human genome according to the reference sequence of the hg19 human assembly. Only DREs in open chromatin, i.e., DREs overlapping the DNase-seq broadPeaks from an ENCODE experiment for a given cell line, were considered for training. Additionally, bound DREs in closed chromatin were also considered for testing purposes. Secondly, we obtained the AhR ChIP-seq bed and bigwig files from ChIP-Atlas (Oki *et al*., 2018) where the original sequencing files have been processed uniformly following a standard processing pipeline. Originally, AhR ChIP-seq data was generated in independent experiments - 1) AhR and ARNT ChIP-seq of MCF-7 cells treated with 10 nM TCDD for 45 minutes from (Lo and Matthews, 2012) - the binding data was obtained from GEO Datasets – accession GSE41820, 2) AhR and AhRR ChIP-seq of MCF-7 cells treated with 10 nM TCDD for 45 minutes and 24 hours from (Yang *et al*., 2018) - the binding data was obtained from GEO Datasets – accession GSE90550, 3) AhR ChIP-chip of T-47D cells treated with 1 µM 3-MC or 10 nM TCDD for 1 hour from (Ahmed *et al*., 2009; Pansoy *et al*., 2010) - the binding data was obtained from their respective publications and converted from hg18 to hg19 using the liftOver tool (Navarro Gonzalez *et al*., 2021) 4) AhR ChIP-seq of primary hepatocytes treated with 1nM of TCDD for 24 hours (this publication) – accessible through GEO Datasets – accession GSEXXXXX; 5) AhR (3xFLAG tagged AhR) ChIP-seq of untreated HepG2 cells from ENCODE – accession (Dunham *et al*., 2012; Davis *et al*., 2018); 6) AhR ChIP-seq of GM17212 cells treated with 1 µM 3-MC for 24 hours - binding data obtained from ChIP-Atlas – accessions SRX4342282, SRX4342283, SRX4342285, and SRX4342286. Details of these experiments are summarized in **Supplementary Table 1** and data sources are summarized in **Supplementary Tables 3 and 4**.

Bound DREs for the purposes of model training were determined as DREs found in open chromatin and under AhR peaks where only one DRE was present under the AhR peak (referred to as singleton DREs). Unbound DREs are DREs in open chromatin found at least 500 base pairs away from the boundary of any AhR peak, as well as 100 base pairs away from any other DRE, in order to minimize confounding of DRE contribution to binding. All other DREs in open chromatin were considered ambiguous and were not used in model training.

### Promoters and enhancers

We obtained all annotated transcription start sites from Ensembl 105 BioMart (human genes; GRCh38.p13) (Howe *et al*., 2021) and considered regions ±200 and ±1500 bp around the TSS as stringent and relaxed promoters, respectively.

We obtained all computationally predicted enhancers from ChromHMM (Ernst and Kellis, 2012) for samples that had ChromHMM data available – HepG2 and GM12878 (ENCODE) and MCF-7 (GEO Datatest – accession GSE57498). Both weak and strong enhancers (ChromHMM states 4 through 7) were considered as valid enchancers.

### Sequence and genomic signal features

For each DRE in the human genome, we have obtained the genomic sequence of seven nucleotides 5’ and 3’ from the DRE (5’-GCGTG-3’). These nucleotides were one-hot encoded and used as features in our machine learning models. In total there were around 1.6 million DREs spread across the human genome. However, only a small fraction of them fulfilled the criteria for bound and unbound DREs used in training and testing.

DNase-seq, as well as all available histone mark and transcription factor ChIP-seq genomic signal (bigwig) files were downloaded for MCF-7, T-47D, primary hepatocytes, HepG2, GM12878 (as the closest match to GM17212 where AhR was ChIP-ed) from the ENCODE consortium. List of all used experiments and files is provided in **Supplementary Table X**. For each bound and unbound DRE and each genomic signal (bigwig) file we extracted the value of the genomic signal 740 base pairs up and downstream from the DRE, for a total of 1495 base pairs of signal (DRE width is 5 base pairs). The extracted signal is split into 15 bins of equal (99 base pairs) size and the signal within each bin is averaged to produce 15 features corresponding to the particular DRE-genomic signal combination. During averaging, any areas of missing signal are replaced with zeros.

### Model architecture and training

For each cell line and all the bound and unbound DREs appearing in open chromatin of that particular cell line, we have created sequence, as well as genomic signal features for all available DNase-seq, histone mark and transcription factor (TF) ChIP-seq. We have then performed hyperparameter tuning of an XGBoost model through a grid search of the hyperparameter space with the following values - max_depth = {3, 4, 5, 6, 7}, min_child_weight = {3, 4, 5, 6, 7}, subsample = {1.0, 0.9, 0.8, 0.7}, colsample_by_tree = {1.0, 0.9, 0.8, 0.7} and eta = {0.05, 0.075, 0.1, 0.125, 0.15, 0.2, 0.3}. We have reported the average performances over all five folds for the best performing models in terms of hyperparameter selection.

### Model evaluation

In addition to evaluating the models through 5-fold cross validation, we also evaluated model performance on predicting the binding status of DREs that occurred under multi-DRE AhR peaks both in open and closed chromatin. For each such peak and each DRE under the peak we used the AhR binding prediction model to make a prediction regarding whether the DRE is bound or not. If at least one DRE under the peak was predicted as bound, the peak was considered recovered, and the total fraction of recovered peaks was reported.

Similarly, we evaluated the model performance on predicting the binding status of AhR peaks without DREs. Briefly, for each 0-DRE peak we simulated five dummy DREs. These dummy DREs are not actually present in the genomic sequence and only represent the genomic location that was used as reference for the calculation of all non-sequence model input features, The center of the first dummy DRE is aligned to the center of the AhR peak and the other four dummy DREs are positioned -100, -50, +50, +100 base pairs relative to the center point of the first dummy DRE. As with the recovery of multi-DRE peaks, a zero-DRE peak is considered partially recovered if at least one of the five dummy DREs is predicted as bound. The peak is considered centrally recovered if the central DRE is predicted as bound.

### Model Performance Metrics

To calculate the area under Receiver Operating Characteristic (auROC) and area under Precision Recall (auPRC) curves we used the output of the XGBoost algorithm in the form of probabilities of each particular observation (DRE) belonging to a particular output class (bound or unbound). By using different thresholds for these probabilities above which the model predicts a DRE as bound, we obtain the number of true and false positives for each threshold, as well as true and false negatives relative to the ground truth of DRE binding obtained from the corresponding AhR ChIP-seq experiment. Each particular threshold produces a point on the ROC and PRC curves; the area under the curve is calculated using a line interpolated through all the points.

### Statistical analysis

Statistical analysis was carried out in Python 3, using the scipy 1.8.0 package (Virtanen *et al*., 2020). ChIP-seq signals were analyzed using the Kruskal-Wallis test and post-hoc analysis performed with the Wilcoxon test for each pair. Results were considered significant if P-value was <0.01.

### Data availability

The primary hepatocyte ChIP-seq sequencing data set generated in this study has been deposited to Gene Expression Omnibus under the accession code GSEXXXXXX.

#### Author contributions

S.B. and S.C. conceived and conceptualized the research. D.F. and W.Q. designed the research, downloaded and processed publicly available data, performed bioinformatic analyses, and created and evaluated machine learning models. O.K. and D.M. helped with the bioinformatics analysis, writing and results interpretation. M.E.A., and E.L.L. oversaw the primary hepatocyte AhR ChIP-seq experiment. S.C., S.B., D.F. and W.Q. co-wrote the manuscript. D.F. generated the figures.Supplementary Tables

## Acknowledgments

This work was partially supported by the USDA National Institute of Food and Agriculture, Michigan AgBioResearch, and the National Institute of Environmental Health Sciences of the National Institutes of Health under awards number R01 ES031937 to S.B and S.C. and P42 ES04911. The content is solely the responsibility of the authors and does not necessarily represent the official views of the National Institutes of Health. This work was supported in part through computational resources and services provided by the Institute for Cyber-Enabled Research at Michigan State University.

## Supplementary Tables

**Supplementary Table 1** – Summary of ChIP-seq experiments (number of peaks, DREs under the peaks, treatment conditions, etc.)

**Supplementary Table 2** – number of AhR peaks found in open chromatin + number of unbound DREs is higher than number of bound DREs

## Supplementary Figures

**Supplementary Figure 1.** Histograms representing DRE position relative to the mid-point of singleton AhR peaks 1) in MCF-7 cells treated with 10 nM TCDD for 45 minutes, 2) not overlapping ARNT peaks in MCF-7 cells treated with 10 nM TCDD for 45 minutes, 3) in primary hepatocytes treated with 1 nM TCDD for 24 hours, 4) in untreated HepG2 cells, 5) in GM17212 cells treated with 1 μM 3-MC for 24 hour.

**Supplementary Figure 2.** Venn diagram representing the overlap of bound DREs in 1) MCF-7 cells treated with 10 nM TCDD for 45 minutes, 2) T-47D cells treated with 10 nM TCDD for 1 hour, 3) T-47D cells treated with 1 μM 3-MC for 1 hour.

**Supplementary Figure 3.** Boxplot of coverage scores in reads per million (RPM) for 0-DRE (blue box), singleton (orange box), multi-DRE (green box) AhR peaks in DMSO (control) treated cells (**left panels**) and AhR agonist (treatment) treated cells (**right panels**) in 1) MCF-7 cells treated with 10 nM TCDD for 45 minutes, 2) primary hepatocytes treated with 1 nM TCDD for 24 hours, 3) untreated HepG2 cells, 4) GM17212 cells treated with 1 μM 3-MC for 24 hours.

**Supplementary Figure 4.** Performance of models predicting the binding status of DREs in open chromatin with five different sets of input features, in 1) untreated HepG2 cells, 2) GM17212 cells treated with 1 μM 3-MC for 24 hours. Performance of each set of features is represented as a mean line with a 95% confidence interval shaded around the line resulting from 5-fold cross validation. The legend shows the list of features used, as well as area under the curve. Both receiver operating characteristic (ROC) - left panel, and precision recall (PRC) curves – right panel, are shown.

**Supplementary Figure 5.** Number of DRE clusters containing N DREs (N = 2, 3,…, 19). A DRE cluster is defined as a 500 base pair wide region containing at least two DREs.

**Supplementary Figure 6.** Feature importance of all local chromatin context features excluding DNA sequence flanking the DRE in models predicting the DRE binding status of DREs in open chromatin of enhancers only. Feature importance is measured as feature importance gain in XGBoost classifier model trained on a particular cell line or type. Feature importance for each bin of a particular chromatin context feature is normalized to the bin with the highest feature importance for that chromatin context feature.

## Notes

### Competing Interest Statement

The authors have declared no competing interest.

