## Supplementary Figures for "Predictive Models of Genome-wide Aryl Hydrocarbon Receptor DNA Binding Reveal Tissue Specific Binding Determinants"

### Slide 1
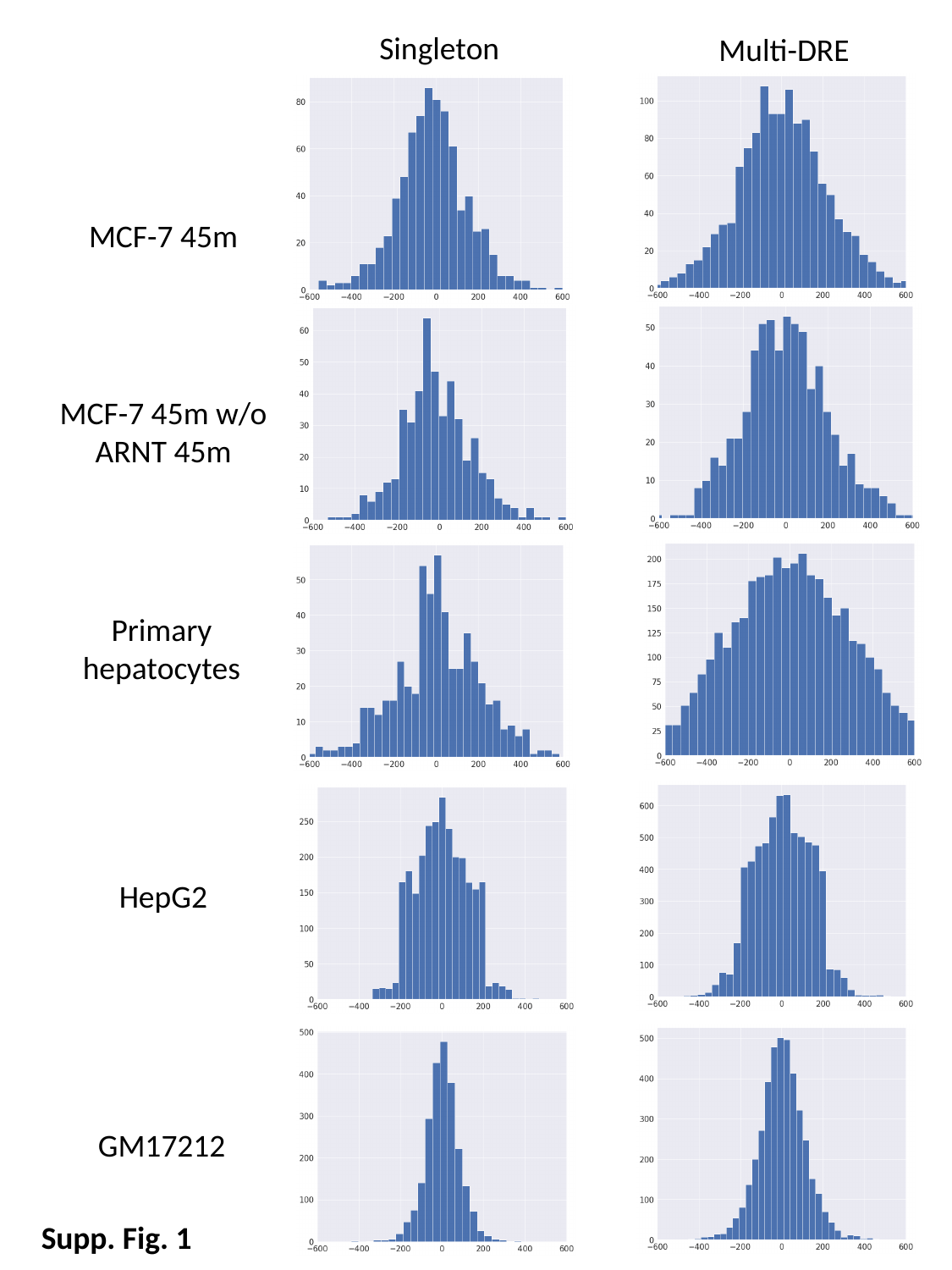

Singleton
Multi-DRE
MCF-7 45m
MCF-7 45m w/o ARNT 45m
Primary hepatocytes
HepG2
GM17212
Supp. Fig. 1

### Slide 2
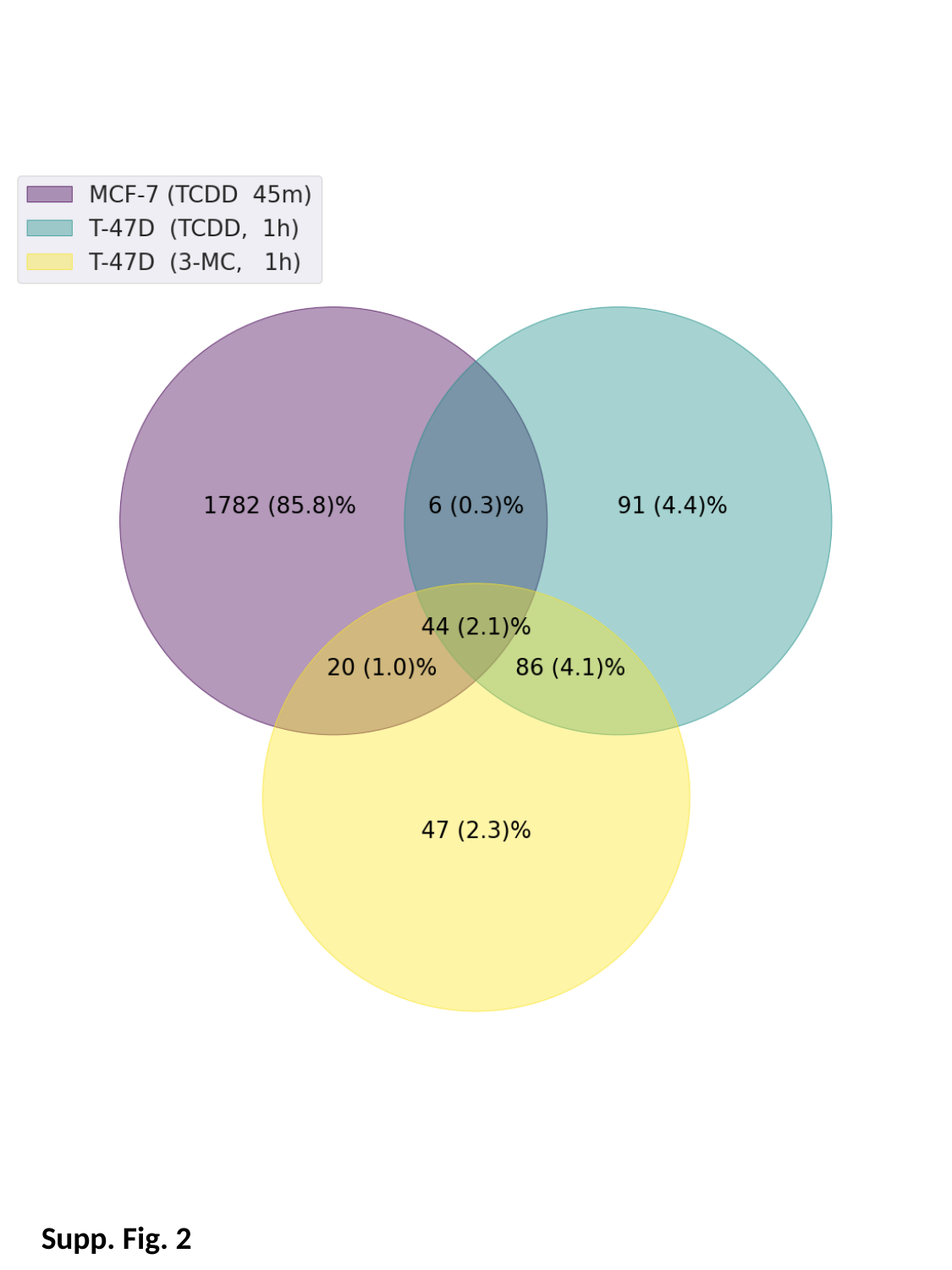

Supp. Fig. 2

### Slide 3
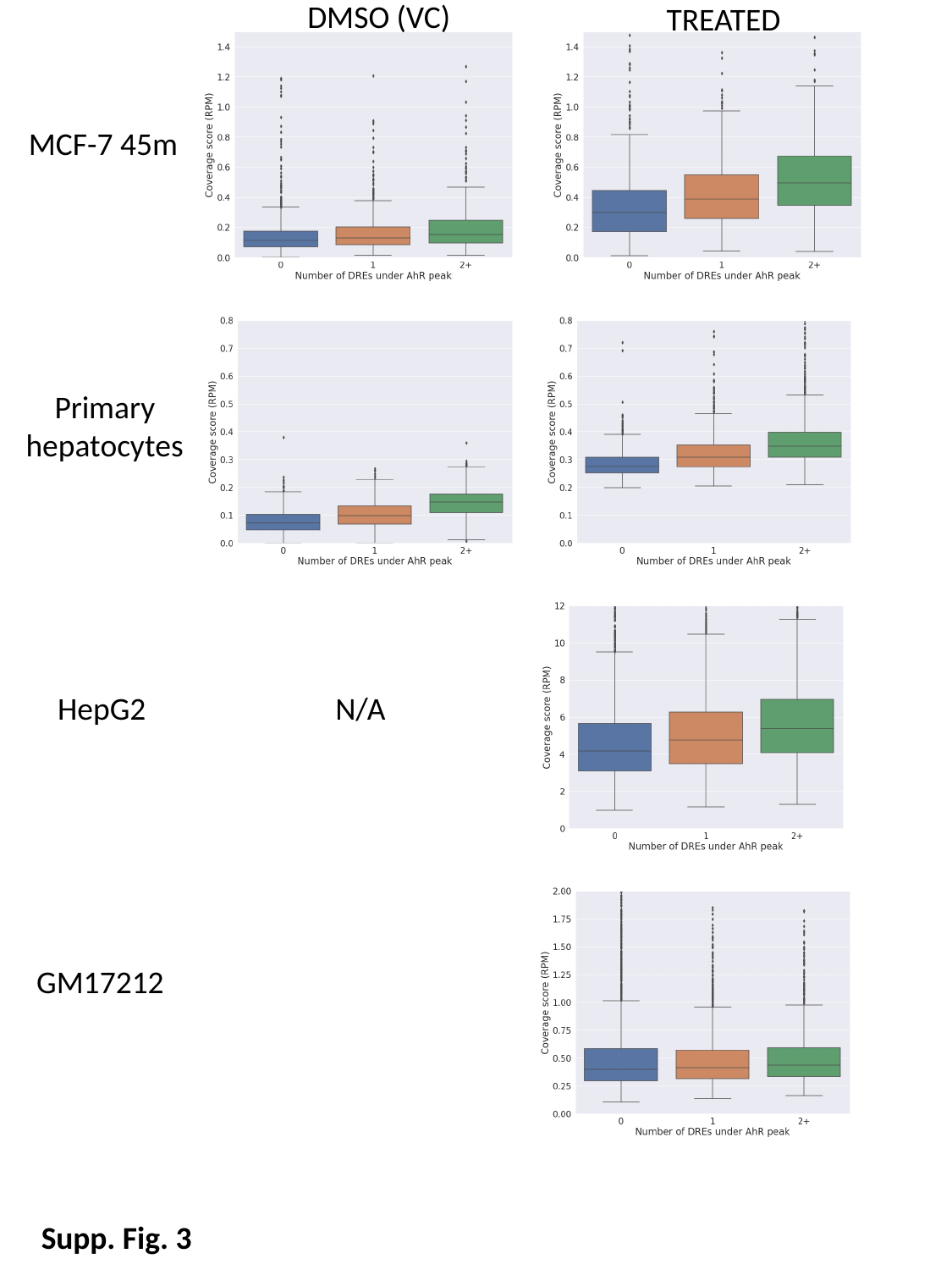

DMSO (VC)
TREATED
MCF-7 45m
Primary hepatocytes
HepG2
N/A
GM17212
Supp. Fig. 3

### Slide 4
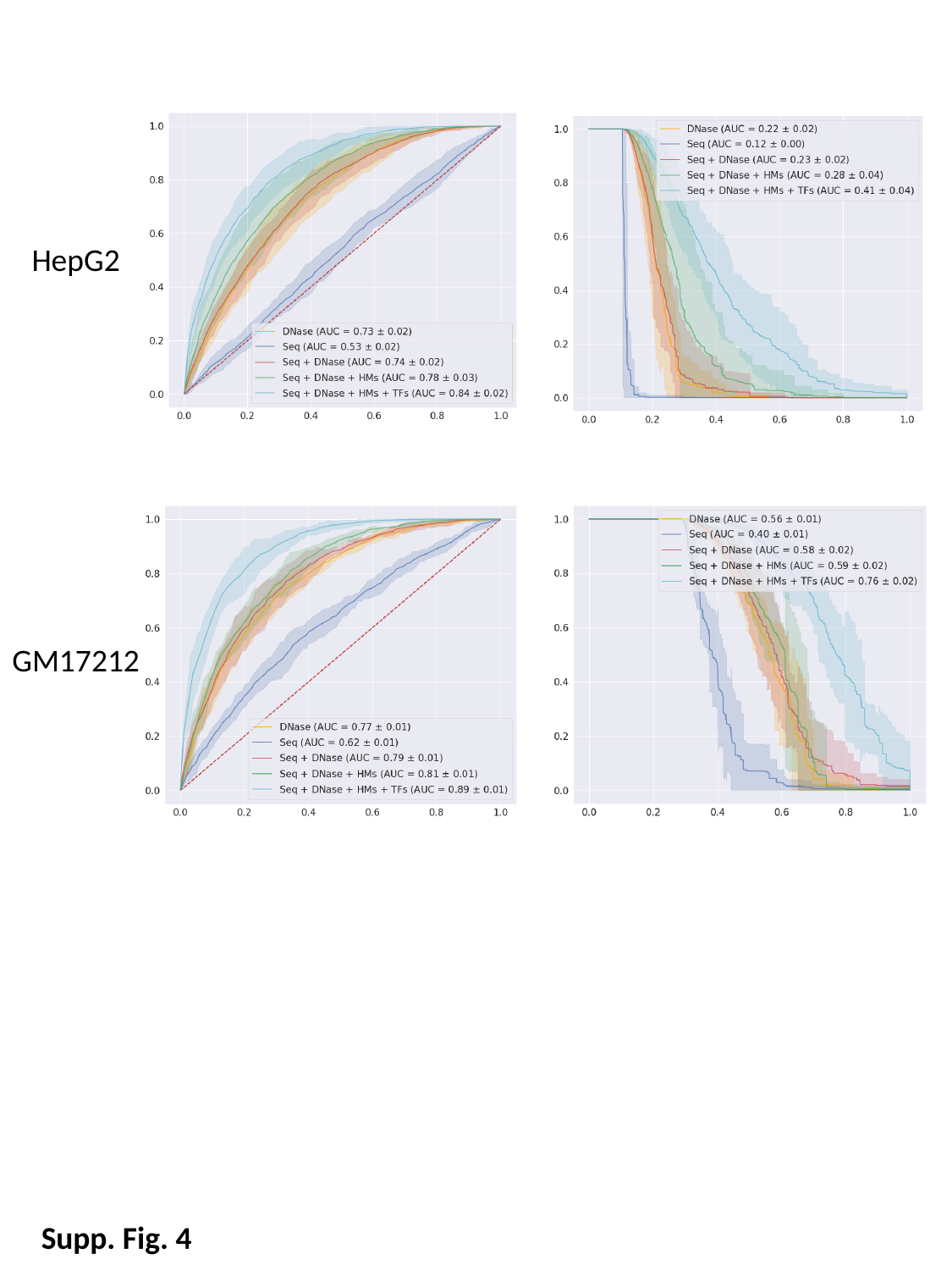

HepG2
GM17212
Supp. Fig. 4

### Slide 5
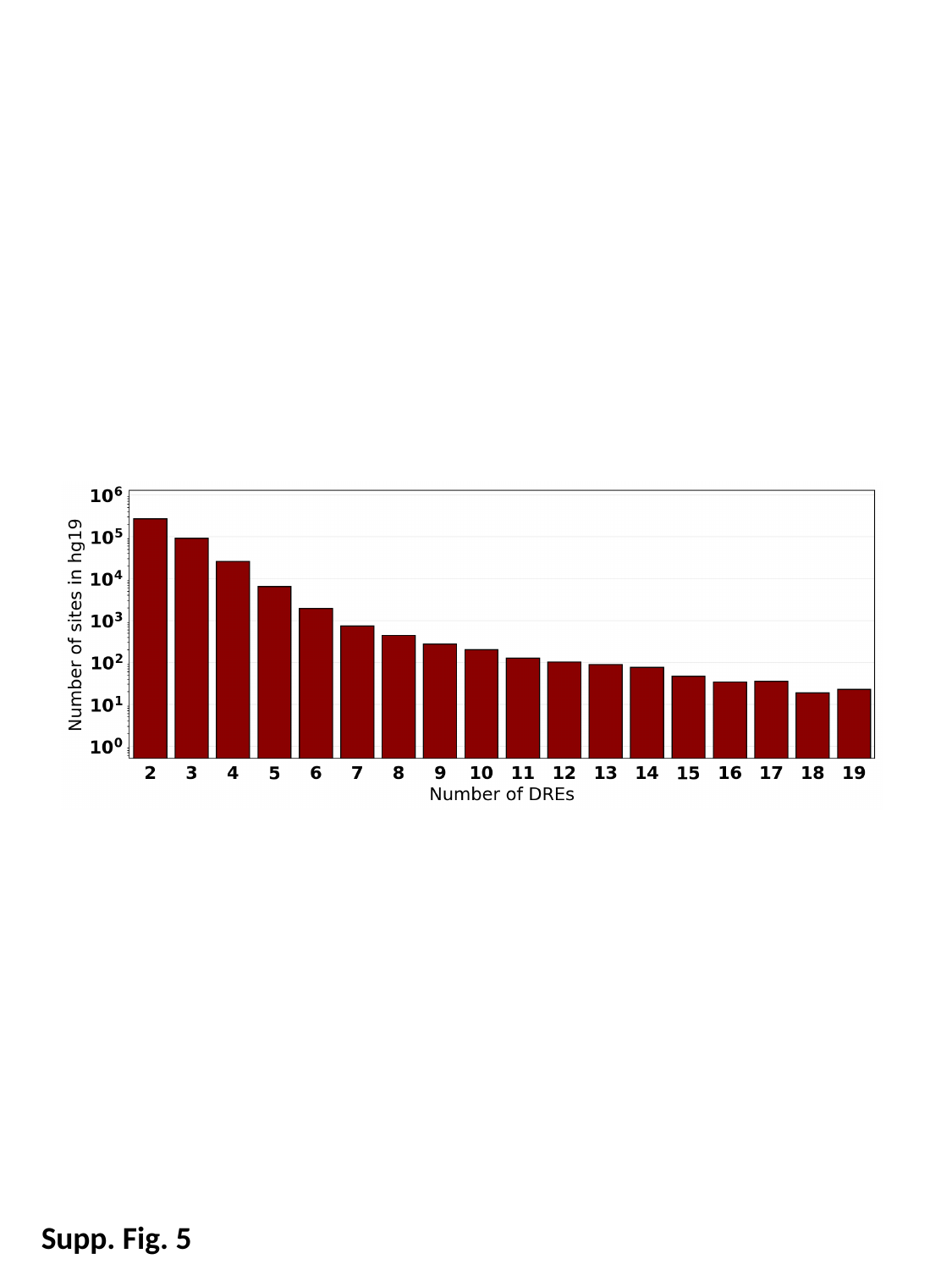

Supp. Fig. 5

### Slide 6
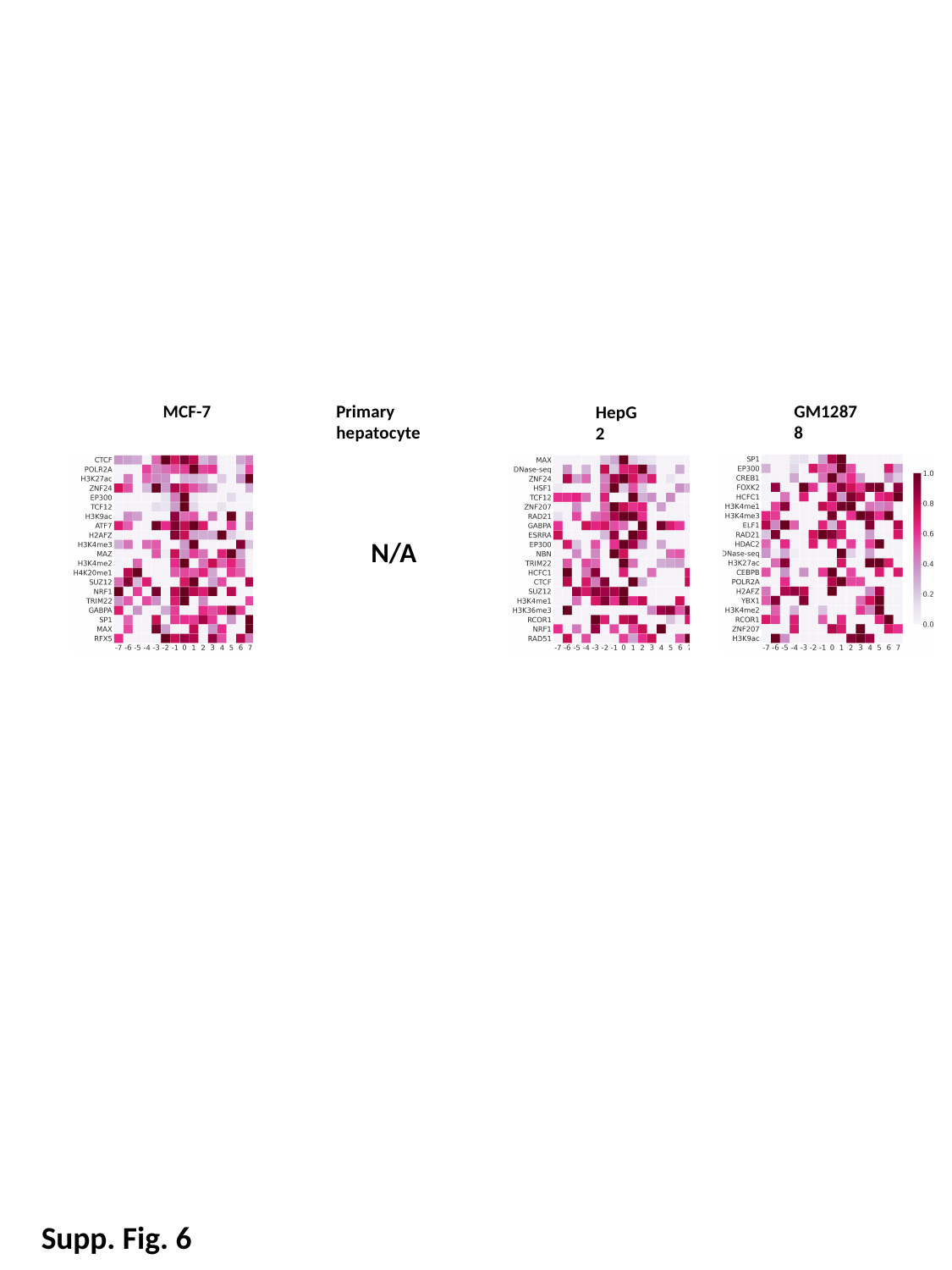

MCF-7
Primary hepatocyte
GM12878
HepG2
N/A
Supp. Fig. 6
